# Synaptopodin is required for stress fiber and contractomere assembly at the epithelial junction

**DOI:** 10.1101/2020.12.30.424702

**Authors:** Timothy Morris, Eva Sue, Caleb Geniesse, William M Brieher, Vivian W Tang

## Abstract

The apical junction of epithelial cells can generate force to control cell geometry and perform contractile processes while maintaining barrier function and adhesion. Yet, the structural basis for force generation at the apical junction is not fully understood. Here, we describe 2 synaptopodin-dependent actomyosin structures that are spatially, temporally, and structurally distinct. The first structure is formed by retrograde flow of synaptopodin initiated at the apical junction, creating sarcomeric stress fibers that lie parallel to the junction and insert into junctional complexes on the apical plane. Retrograde flow of synaptopodin is also seen at vinculin-decorated basal junctions on the basal plane. Contractions of apical stress fibers is associated with clustering of membrane complexes via side-on synaptopodin linkers whereas contractions of stress fibers inserted at the apical junction via head-on synaptopodin linkers results in junction shortening. Upon junction maturation, apical stress fibers are disassembled. In mature epithelial monolayer, a motorized “contractomere” complex capable of “walking the junction” is formed at junction vertices. Contractomere motility results in changes in junctional length, altering the overall shape of the cell and packing geometry within the monolayer. We propose a model of epithelial homeostasis that utilizes contractomere motility to preserve the permeability barrier during intercellular movement and junctional processes.

**Summary Statement:** Synaptopodin retrograde flows initiate the assembly of apical and basal stress fibers from the apical and basal junctions. In mature apical junction, a motorized junctional complex, we termed the contractomere, allows the apical junction to change length and organize cell geometry within a confluent monolayer.

## Introduction

Epithelial cells cover body cavities and line internal organs, providing continuous protection from biological, chemical, and mechanical insults. The maintenance, stability, and physiological functions of an epithelium require the assembly of a specialized cell-cell junction known as the apical junction. Epithelial apical junction provides adhesions between cells, forms permeability barrier against macromolecules and microorganisms, and regulate epithelial cell packing and geometry. Force exerted by the epithelial junction not only can reshape cell boundaries and adjust junctional length but also facilitate cell rearrangement during dynamics processes including wound migration and morphogenesis (Pinheiro and Bellaiche, 2018). Moreover, junction contractility is necessary for cell extrusion and purse-string wound closure that are essential for epithelial homeostasis (Bement et al., 1993; Clark et al., 2009; Danjo and Gipson, 1998; Florian et al., 2002).

Force generation at the junction depends on myosin II, a >300 nm bipolar minifilament consisted of anti-parallel arrangement of barbed-end-directed myosin II motors. Activation of junctional contractility results in shortening of the junction and destabilizes E-cadherin adhesions (Cavanaugh et al., 2020). In contrast, application of orthogonal force to the junction results in strengthening of E-cadherin adhesions (Gomez et al., 2011; Kannan and Tang, 2015; Kannan and Tang, 2018; le Duc et al., 2010). Thus, the biological outcome of force at the junction appears to depend on the orientation of the applied force, which is the directional information embedded within the force vector.

What is the molecular and structural basis that allows the generation of orthogonal and parallel force at the apical junction? How does the cell organize actomyosin populations to control the direction of force? Can the junction pull on actin filaments or is the junction being pulled by actomyosin? Do we know whether contractile force is generated at the junction or is the junction on the receiving end of cytoplasmic force? At least 2 actomyosin structures could exert orthogonal force on the apical junction. The apical-medial actomyosin meshwork in polarized epithelial cells can generate isotropic contractile force, pulling the junction orthogonally (Roper, 2015). Contraction of cytoplasmic actomyosin network also can exert force on cell-cell adhesions (Kannan and Tang, 2015; Wu et al., 2014). Myosin IIA is organized into actin cables underneath the junction whereas myosin IIB is found on membrane adhesions. Contractions of these 2 actomyosin populations could exert parallel force on the junction (Heuze et al., 2019). Earlier studies have shown that myosin IIB controls actin accumulation whereas myosin IIA regulates E-cadherin stability (Smutny et al., 2010). Furthermore, myosin IIB, but not myosin IIA, plays a role in alpha-catenin mechanoregulation (Heuze et al., 2019). Thus, different myosin II isoforms can form actomyosin structures to serve distinct force-dependent functions.

In this paper, we describe 2 novel actomyosin structures, apical stress fiber and a motorized organelle we name “contractomere”. We provide evidence for an essential molecular component, synaptopodin, for their assembly.

## Results

To understand how actomyosin organization contributes to force generation at the epithelial junction, we focused on 3 actin-binding proteins, alpha-actinin-4, synaptopodin, and myosin IIB, which are known regulators of cellular contractility (Asanuma et al., 2005; Hotulainen and Lappalainen, 2006; Kannan and Tang, 2015; Mundel et al., 1997; Solinet and Vitale, 2008). Synaptopodin is a vertebrate-specific protein expressed ubiquitously in human (Uhlen et al., 2015). Using antibodies against different spliced regions, we showed that epithelial and endothelial cells express different variants of synaptopodin (Fig S1A). Synaptopodin decorates the apical junction and stress fibers of epithelial and endothelial cells (Fig S1B). In MDCK cells, synaptopodin, alpha-actinin-4, and myosin IIB form a network of stress fibers connecting cell-cell adhesions across multiple cells (Fig S2A-B). Thin-section electron microscopy of T84 intestinal epithelial cells revealed that apical stress fiber with distinctive “sarcomere” is attached via actin thin-filaments to the apical junction (Fig S2C). In this paper, we focus on synaptopodin actomyosin structures in MDCK cells.

### Organization of 2 novel actomyosin structures at the apical junction

MDCK cell monolayers grown on Transwells develop apical junctions with strong cell-cell adhesion and a robust permeability barrier (Kannan and Tang, 2015; Tang and Goodenough, 2003). We grew MDCK cells on Transwells and compared their actomyosin structures at various stages of junction development using superresolution microscopy. At an early stage of junction development, myosin IIB and synaptopodin exhibit periodic and alternating pattern (Fig 1A, S1B & S2A). Apical stress fibers are positioned parallel to the junction and spatially separated from E-cadherin junctions except at points of insertion (Fig 1A, orange arrowheads). This is analogous to basal stress fibers where attachments at focal adhesions occur at the ends of the stress fibers. Alternating pattern of myosin II and alpha-actinin is a signature organization for contractile stress fibers (Naumanen et al., 2008; Pellegrin and Mellor, 2007). During junction development, apical stress fibers form side-on interactions with the apical junction via synaptopodin linkers (Fig 1B, arrowheads). Upon association with E-cadherin junctions, apical stress fibers become less organized, and the characteristic sarcomeric-repeats are less recognizable. As junctions mature over time, the alternating pattern of myosin IIB and synaptopodin completely dissolves (Fig 1C). Concomitantly, E-cadherin, myosin IIB, and synaptopodin accumulate at the ends of linear junctions, also known as junction vertices. Upon maturation of the epithelial monolayer, apical stress fibers are absent, and a novel complex containing myosin IIB, synaptopodin, and alpha-actinin-4 is formed at junction vertices (Fig 1D). We refer to these 2 structures as “type I” when myosin IIB and synaptopodin are arranged in alternating pattern and “type II” when they overlap (Fig 1E). We termed the type II structure “contractomere” since it has not been described before; “contracto” refers to contractile activity of myosin II that can pull on actin filaments during isometric, eccentric, or concentric contraction; “mere” refers to it being a standalone structure that can be isolated biochemically. The goal of this paper is to determine whether these actomyosin assemblies are associated with junctional processes and whether synaptopodin is required for their assembly.

**Figure 1.**
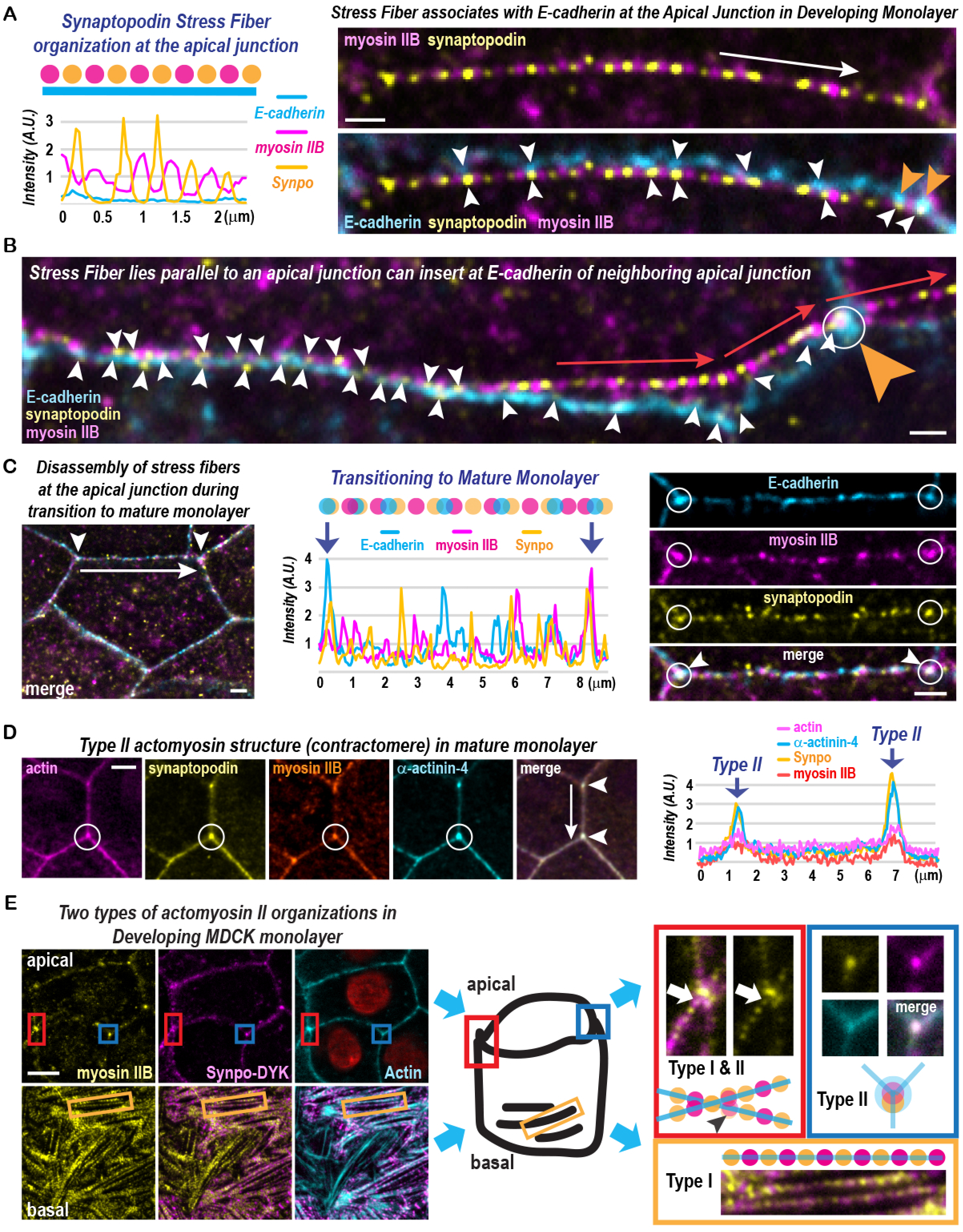
Superresolution microscopy of 2 actomyosin structures at the apical junction. (A) Apical stress fiber has alternating myosin IIB and synaptopodin pattern and lies parallel to E-cadherin junction. Orange arrowheads point to the end of an apical stress fiber marked by E-cadherin. White arrowheads point to the junctional region where synaptopodin is in proximity to E-cadherin. White long arrow shows the junctional length used for x-axis of the graph. Scale bar is 500 nm. (B) Apical stress fiber is inserted head-on at E-cadherin junction, marked by orange arrowhead. Synaptopodin overlaps with E-cadherin at the insertion point of apical stress fiber, in white circle. White arrowheads mark sites where synaptopodin is in proximity to or overlapping with E-cadherin. Scale bar is 500 nm. (C) Disassembly of apical stress fibers in maturing junction results in the loss of alternating synaptopodin and myosin IIB pattern. Disassembly of apical stress fiber coincides with the formation of type II actomyosin structures containing myosin IIB, synaptopodin, and alpha-actinin-4, marked by arrows in graph. White circles in the left panels show type II structures. White arrowheads mark the colocalization of myosin IIB, synaptopodin, alpha-actinin-4 in right panels. White long arrow shows the junctional length used for x-axis of the graph. Scale bars are 1 micron. (D) Apical stress fibers disappear upon junction maturation whereas type II actomyosin structures are prominent at mature apical junctions. Actin accumulates at type II actomyosin structure, circled in white, colocalizing with myosin IIB, synaptopodin, and alpha-actinin-4, as marked by white arrowheads. Graph shows the absence of apical stress fiber and the presence type II structure, named contractomere. White long arrow shows the junctional length used for x-axis of the graph. Scale bar is 1 micron. (E) Two actomyosin structures at the apical junction of MDCK cells. Apical stress fibers are labelled as Type I actomyosin structure with alternating synaptopodin and myosin IIB organization. In maturing junction, Type I apical stress fiber coexists with Type II contractomeres with overlapping synaptopodin and myosin IIB. Basal stress fibers have alternating synaptopodin and myosin IIB organization, same as Type I apical stress fibers. Scale bar is 2 microns.

### Synaptopodin stress fibers are inserted at both apical and basal junctions

Synaptopodin has emerged as one of the best markers for actin cytoskeletal structures. To image stress fiber dynamics at the apical junction, we co-expressed synaptopodin with ZO-1 (Fig 2A & S3A), which is a junction protein that interacts with synaptopodin and alpha-actinin-4 (Chen et al., 2006; Van Itallie et al., 2013). ZO-1 has previously been shown to decorate apical and basal junctions (Anderson et al., 1988; Danjo and Gipson, 1998; Millan et al., 2010; Stevenson et al., 1988; Tornavaca et al., 2015). In MDCK monolayers, synaptopodin apical stress fibers (Fig 2A, blue boxe) are spatially segregated from synaptopodin basal stress fibers (Fig 2A, yellow boxe). Both apical and basal stress fibers can attach to the junction via end-on interactions (Fig 2A & S3A). When an apical stress fiber is attached head-on, the junction would experience orthogonal force. Live-imaging shows that contractions of end-on apical stress fibers are associated with junction shortening (Fig 2B-C, movie 1) and cell shape changes (Fig 2D, movie 2).

**Figure 2.**
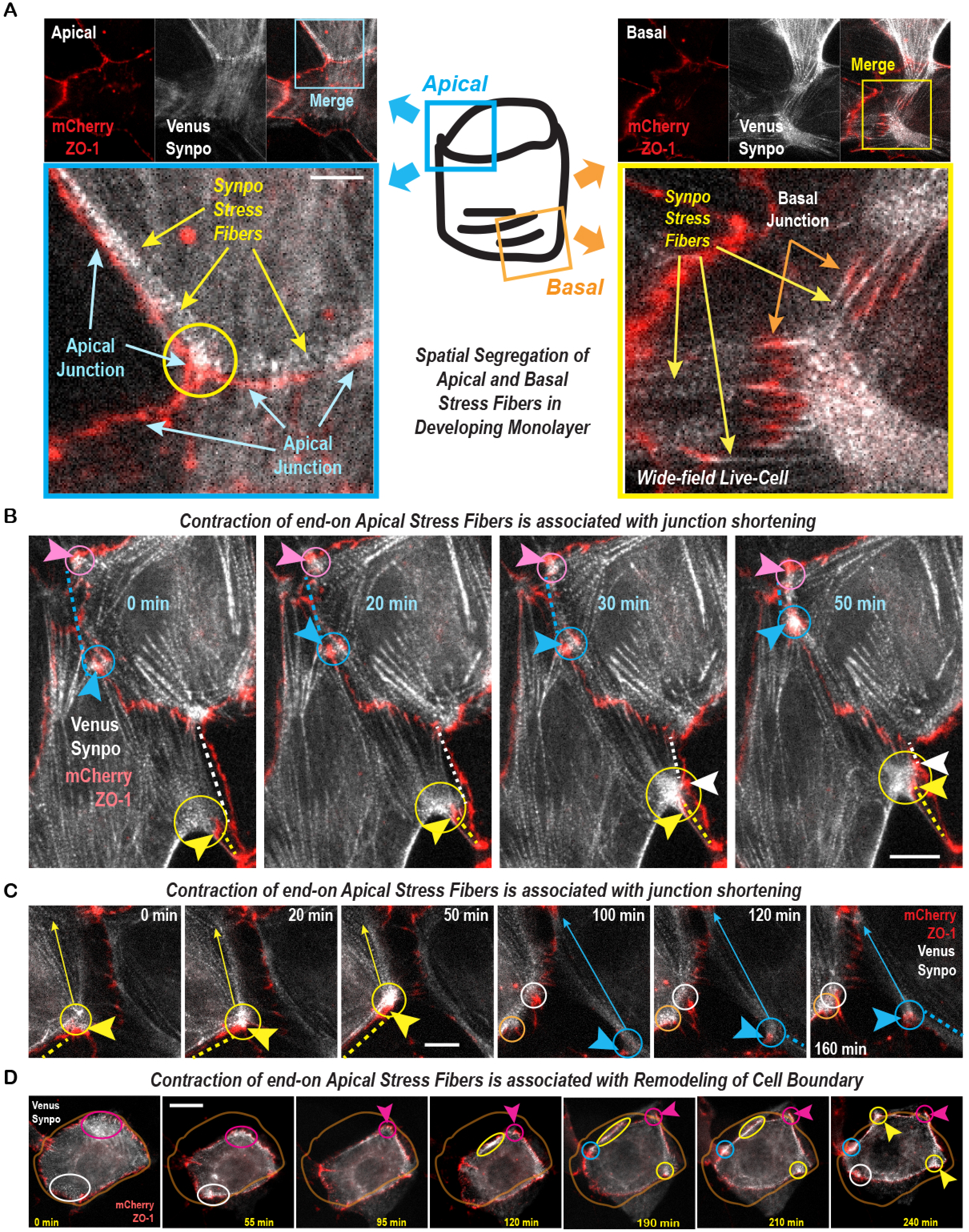
Synaptopodin apical stress fibers are contractile structures distinct from basal stress fibers. (A) Frames taken from a time-lapse of synaptopodin and ZO-1 showing apical and basal stress fibers on different focal planes. Apical and basal stress fibers are inserted at apical and basal ZO-1 junctions, respectively. Scale bar is 1 micron. (B) Frames taken from a time-lapse of synaptopodin and ZO-1. Arrowheads point to sites of stress fiber attachment at the apical junction. Contraction events are circled. Dotted lines represent lengths of linear junction. Scale bar is 5 micron. (C) Frames taken from a time-lapse of synaptopodin and ZO-1. Arrowheads point to sites of stress fiber attachment at the apical junction. Contraction events are circled. Arrows point to direction of movement of the attached junction. Scale bar is 1 micron. (D) Frames taken from time-lapse of synaptopodin and ZO-1. Outline of a cell at time zero is drawn in all panels. Arrowheads point to stress fiber attachment sites at the apical junction. Contraction of apical stress fiber is circled. Scale bar is 10 microns.

We also imaged synaptopodin with vinculin, a junction marker in epithelial and endothelial cells (Efimova and Svitkina, 2018; Huveneers et al., 2012; Kobielak et al., 2004; le Duc et al., 2010; Schnabel et al., 1990; Tornavaca et al., 2015; Twiss et al., 2012; Zhang et al., 2005). In sparsely-plated MDCK cells, synaptopodin stress fibers are inserted end-on into vinculin-decorated basal junctions and focal adhesions on the basal focal plane (Fig S3B).

### Retrograde flow of synaptopodin during stress fiber assembly

In developing monolayer, synaptopodin is continuously incorporated at basal junctions, creating a retrograde flow of synaptopodin along the long axis of basal stress fibers (Fig S4A-C & movie 3-5). In polarized cells, basal stress fibers are several microns away from apical stress fibers on a different focal plane (Fig S4D). On the apical plane, retrograde flow of synaptopodin is initiated at the apical junction (Fig S5A, movies 6-8). In contrast to the basal junction, synaptopodin retrograde flow from the apical junction give rise to stress fibers that are parallel to the junction; this is a 90-degree change in orientation of the stress fibers with respect to the direction of synaptopodin flow (Fig 3A & movie 9), implying distinct mechanisms for stress fiber assembly at the apical and basal junctions.

**Figure 3.**
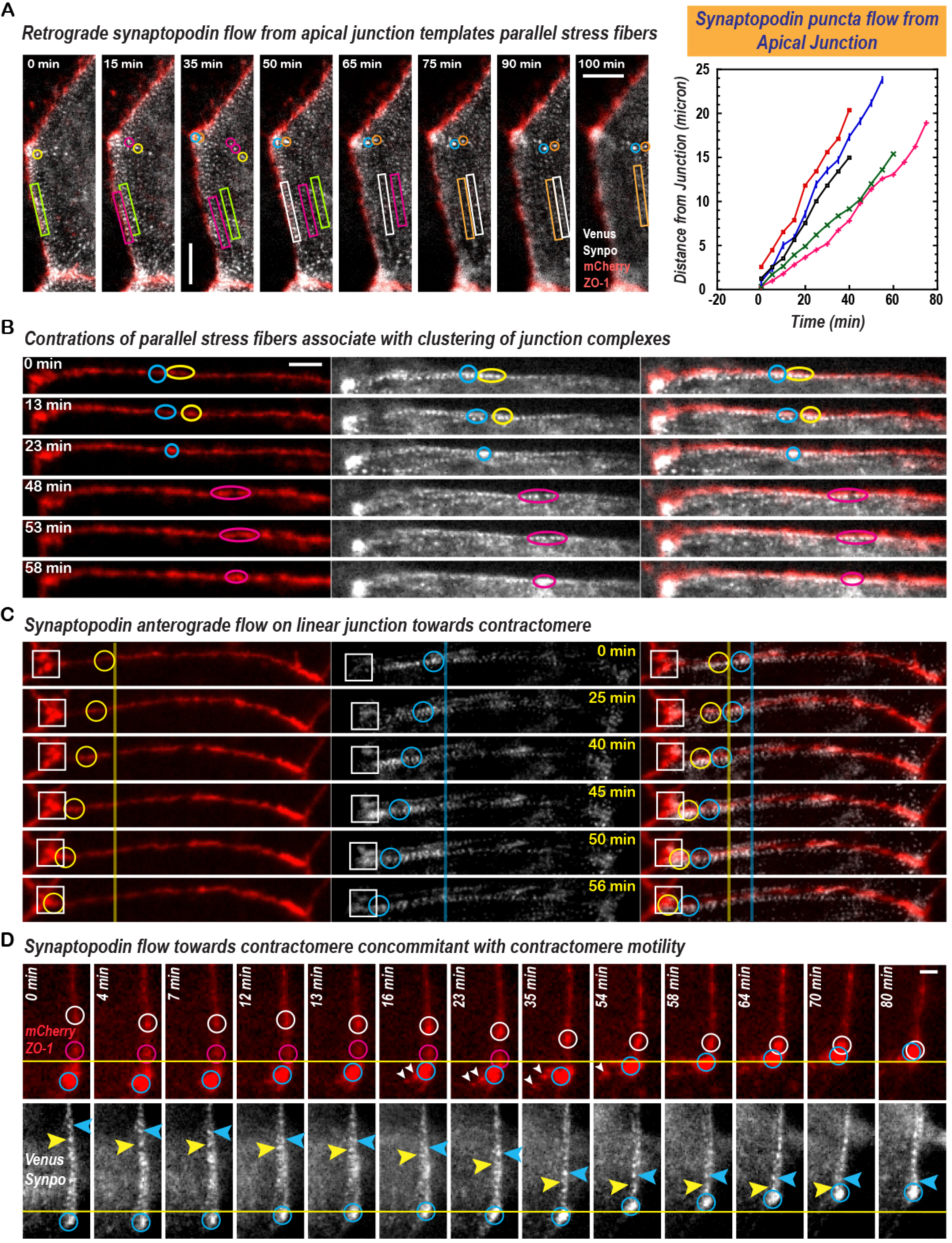
Synaptopodin flow and the evolution of apical stress fibers during junction development. (A) Frames from time-lapse of synaptopodin and ZO-1 in developing junctions. Retrograde synaptopodin flow originates from the apical junction. Rectangles track a row of synaptopodin densities as they flow inward away from the apical junction. Circles track single synaptopodin densities. Graph shows the rate of synaptopodin flow by tracking individual synaptopodin densities. Scale bar is 2 microns. (B) Frames from time-lapse of synaptopodin and ZO-1 of maturing junctions. ZO-1 densities, marked by arrowheads, are temporarily clustered when synaptopodin stress fiber contract (circled). Scale bar is 1 micron. (C) Frames from time-lapse of synaptopodin and ZO-1 at late stage of junction maturation. Anterograde synaptopodin flow towards junction vertex is associated with the movement of vertex against the direction of synaptopodin flow. Circles track synaptopodin densities flowing into the junction vertex. Squares track the movement of junction vertex. Scale bar is 500 nm. (D) Frames from time-lapse of synaptopodin and ZO-1 at late stage of junction maturation. Synaptopodin flowing toward junction vertex is associated with movement of the vertex against the direction of synaptopodin flow. Arrowheads track synaptopodin densities flowing into junction vertex. Blue circles track the movement of junction vertex. Red and white circles track ZO-1 densities moving toward the vertex. Scale bar is 500 nm.

### Contractility and evolution of apical stress fibers during junction maturation

As the junction develop over the next few days, retrograde synaptopodin flow ceases. Synaptopodin apical stress fibers now exhibit contractility, resulting in local clustering of junctional components (Fig 3B & S5B, movies 10-13). In the following few days, synaptopodin apical stress fibers begin to flow towards junction vertices (Fig 3C & movies 14-15). At the vertices, nascent contractomere move against the direction of synaptopodin flow, implicating an energy-driven process is at play at the contractomere (Fig 3D). Motility of a contractomere can simultaneously shorten a junction while lengthening an adjacent junction (Fig S5C & movie 17). Two contractomeres flanking a junction can shorten the junction by moving towards each other (movie 16). This behavior coincides with the dissolution of synaptopodin stress fibers and the assembly of contractomeres in mature monolayer (Fig S6A-B), a type II structure that we described in Fig 1. In developing and mature monolayers, vinculin is absent from the apical junction (Fig S6C-D).

### Contractomere motility in developing monolayer

To understand the relationship between contractomere motility and cell geometry, we performed live-cell structured-illumination microscopy (Fig 4A). In maturing monolayers, individual cells move among each other while maintaining the same neighbors (Fig 4A, lower panels). By tracking individual contractomeres, we found no directional bias in the movement of the contractomeres (Fig 4A, top panels). Measurement of junction lengths showed that contractomere movement can result in either shortening or lengthening of a junction (Fig 4B). Junction shortening occurs when 2 contractomeres move towards each other whereas junction lengthening occurs when 2 contractomeres move away from each other (Fig 4C). These findings indicate that the lengths of individual junctions, the overall shape of the cell, and the packing geometry of the epithelium may be adjusted quickly by sliding contractomeres around the apical cell boundary of in a confluent monolayer of cells. If this hypothesis is true, junctional lengths of a given individual cell would stay constant. We assessed our hypothesis by culturing a confluent monolayer of MDCK cells on collagen I substrate to promote intercellular movement. Under this condition, MDCK cells would exchange neighbors while maintaining constant cell perimeters (Fig 4D). This result supports our hypothesis that contractomere motility can adjust junction proportion and plays a role in regulation of cell geometry in a confluent epithelial monolayer.

**Figure 4.**
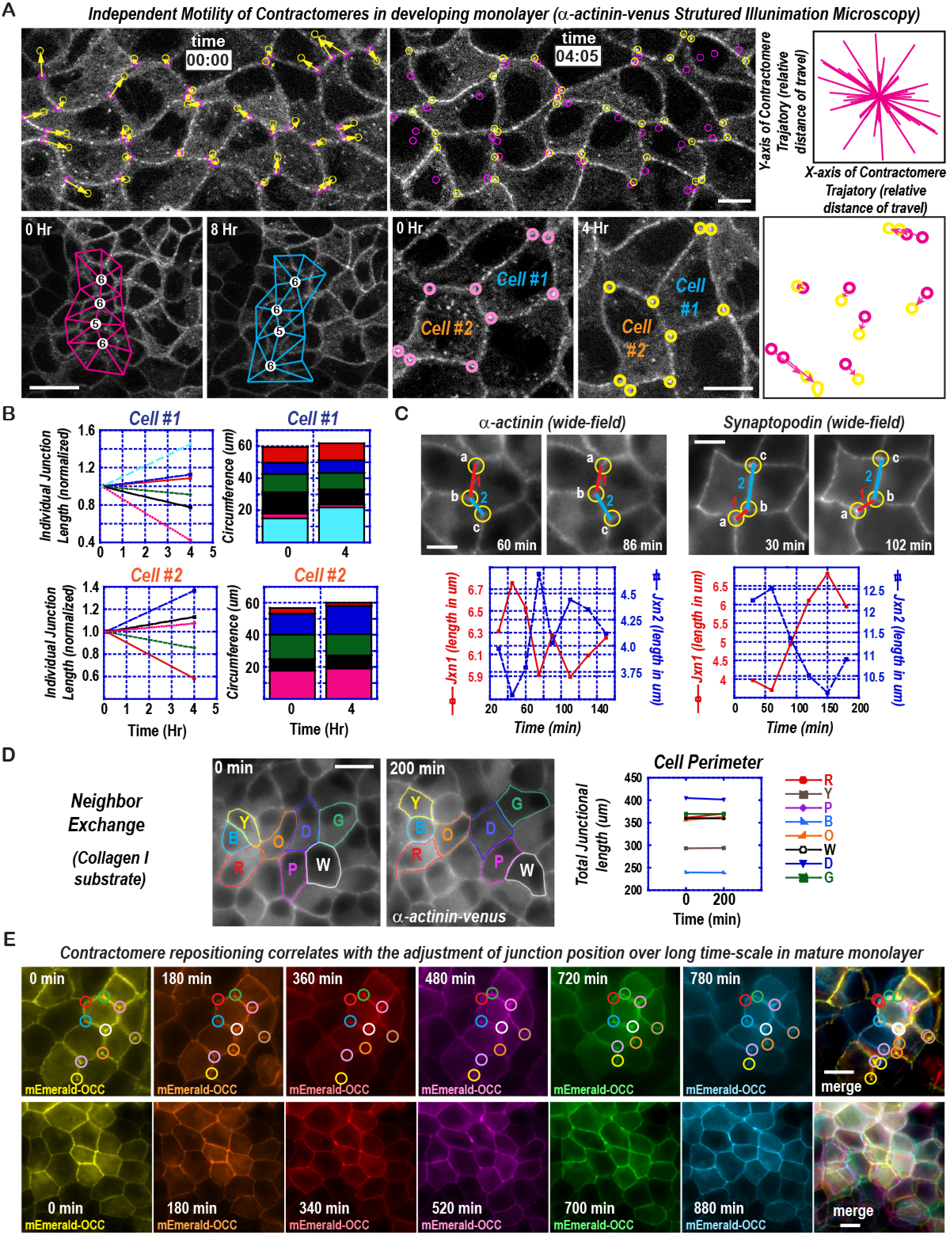
Contractomere motility conserves junctional length. (A) Frames from time-lapse structured-illumination movie of venus-alpha-actinin-1 showing contractomere at the beginning (pink circles) and at the end (yellow circles) of a 4-hour movie. During intercellular organization, junctional length can shorten when 2 contractomeres move towards each other and junctional length can extend when 2 contractomeres move away from each other. The distance and direction of travel for the contractomeres are shown in upper right panel. Lower left 2 panels show the cells maintaining their neighbors during the 4-hour movie. Lower right 3 panels show contractomere motility in 2 cells. Scale bars are 10 microns. (B) Measurement of junctional lengths of the 2 cells in A. Cells #1 and #2 are used to illustrate that some junctions increased but the others decreased their length, resulting in near zero net change in total junctional length. (C) Frames from time-lapse wide-field of venus-alpha-actinin-1 and synaptopodin-venus. Gliding of contractomeres to reproportion junctional lengths. In the a-actinin movie, motility of contractomeres resulted in shortening the junction between contractomeres a and b with concomitant lengthening of the junction between contractomeres b and c. In the synaptopodin movie, motility of contractomeres resulted in lengthening of the junction between a and b with concomitant shortening of the junction between contractomeres b and c. Scale bars are 5 microns. (D) Plating of MDCK cells on collagen I at confluent density resulted in intercellular movement and neighbor exchange. Total junctional length of individual cells remained constant. Scale bar is 10 microns. (E) Frames from time-lapse of occludin showing contractomere and junction movement over 12 hours. Contractomeres are circled.

### Contractomere motility in mature monolayer

Upon epithelial maturation, gross movement of contractomeres subsides. However, at faster movie acquisition rate, contractomeres continue to oscillate at a smaller length-scale and a faster timescale (movies 18-20). This is consistent with the motor property of myosin IIB, which has the ability to step forward and backward, allowing myosin IIB to oscillating on actin filaments (Norstrom et al., 2010). These observations are in agreement with previous study showing junction oscillation can be blocked by blebbistatin (Kannan and Tang, 2015). Beside oscillating back-and-forth at smaller length-scales, junctional lengths and inter-contractomeric distances remain unchanged over long periods of time regardless of individual junctional lengths. Structured-illumination live-imaging showed that apical junctions have curved or wavy contours, suggesting a lack of membrane line tension (Fig S7 & movies 21-24).

### Contractomere motility during cell extrusion

Epithelial monolayer is continuously renewed during homeostasis by balancing cell proliferation and cell death. Removal of unwanted or dying cells are carried out in part by constriction of the apical junction in a process called cell extrusion (Kuipers et al., 2014; Madara, 1990). We imaged alpha-actinin to assess if contractomere motility plays a role in cell geometry organization during cell extrusion. When the apical junction constricts, contractomeres surrounding the extruding cell move towards each other (Fig S8A-B, movies 25-26). Tracking the trajectories of individual contractomeres revealed that new cell-cell interfaces are formed between neighbors of the extruding cell (Fig S8C, blue and orange arrows & movie 27). At the end of junction constriction, contractomeres became very close to each other and the fluorescence signals were overwhelmed by out-of-focus light from the extruding cell. To visualize contractomeres without the interference by out-of-focus light, we imaged cell extrusion using structured-illumination live-cell microscopy (Fig S8D-E & movies 28-29). By track contractomeres during a constriction, we found that contractomere movements can close the apical junction completely during cell extrusion, preserving the continuity of the epithelial cell sheet during the entire process.

To image the apical junction and synaptopodin simultaneously during cell extrusion, we use occludin, a ZO-1-interacting membrane protein. We found that junctional lengths remain relatively unchanged over a long period of time in mature monolayers (Fig 4E). Macroscopically, cell extrusion is not accompanied by gross cell reorganization (Fig S8F & movie 30-31), indicating that it must be achieved through local junctional changes.

Live-cell imaging of synaptopodin and occludin shows that contractomere move during cell extrusion, resulting in elongation of the apical junction (Fig 5A & movie 32). By measuring the distance between a motile and an immobile contractomere, we found that contractomere motility is characterized by persistent “run” periods with intermittent “pause” periods (Fig 5A, graph). As we have seen earlier (Fig 4E & S8F), cells in homeostatic monolayer are relatively stationary, thus could not have contributed to contractomere motility. In homeostatic monolayer, there are 2 populations of contractomeres during cell extrusion, a mobile pool and an immobile pool (Fig 5B & Movies 33-35). While the contractomeres surrounding the extruding cell move to constrict the apical junction (Fig 5B, lower left graph), the rest of contractomeres in the same cells are immobile (Fig 5B, lower middle graph). As the apical junction constrict, the perimeter of the extruding cell decreases; this is concomitant with extension of junctions flanking the constricting contractomeres (Fig 5B, lower right graph). Individual contractomeres have slightly different behaviors with unique extension velocities during “run” periods and “pause” periods (Fig 5B, graphs). By measuring the perimeter of the extruding cell and the lengths of the extending junctions, we found that contractomere motility alone could account for conservation of total junctional length during in the first half of cell extrusion (Fig 5C). This is especially striking when contractomere motility in a mature monolayer would extend the length of the junction without affecting the lengths of other junctions or the positions of other contractomeres (Fig 5D).

**Figure 5.**
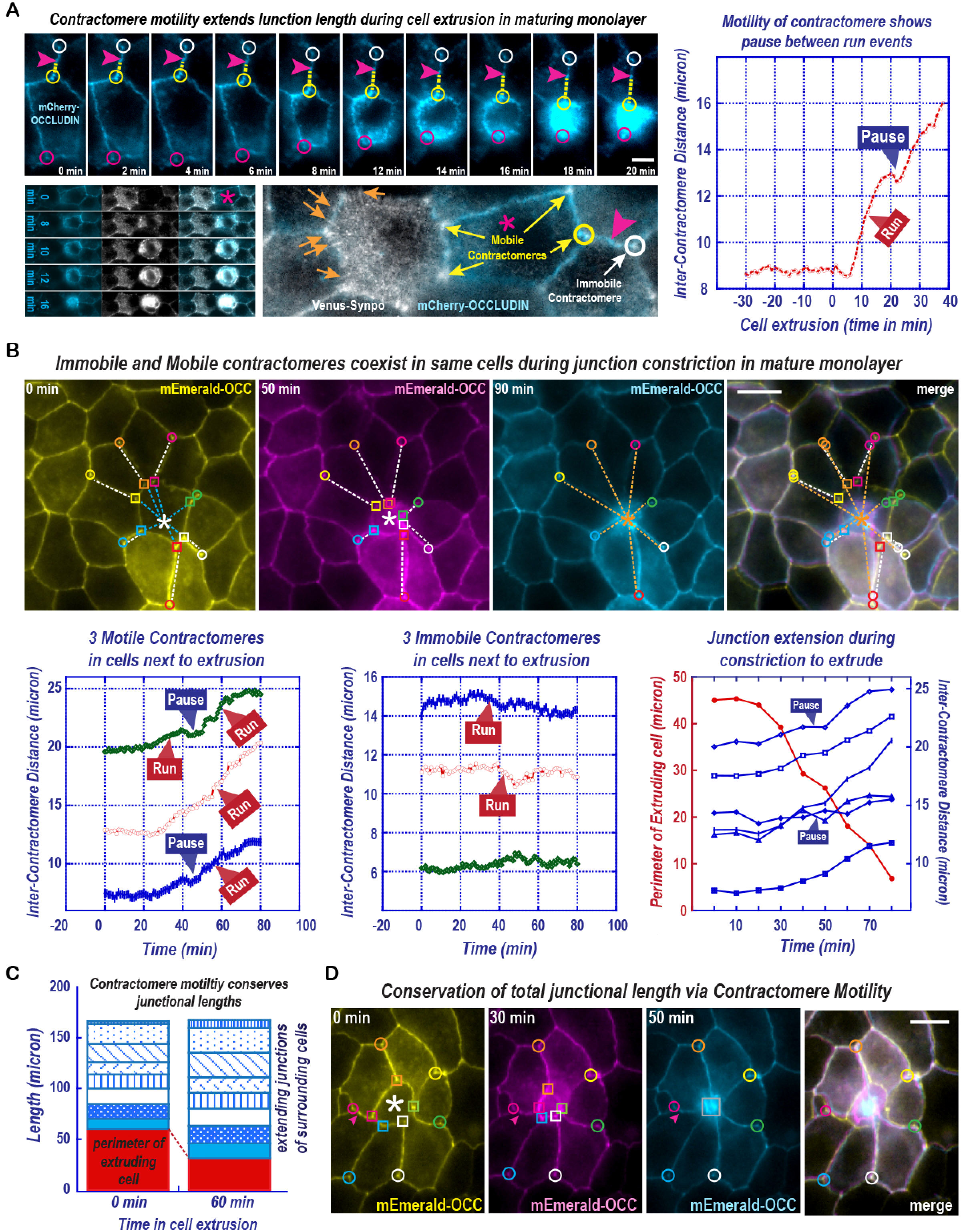
Mobile and immobile contractomeres. (A) Frames from time-lapse showing contractomere motility during cell extrusion in maturing monolayer. Arrowheads track the location of a stationary ZO-1 density. Graph shows inter-contractomere distance between a motile contractomere (yellow circle) and a relatively immobile contractomere (white circle). Scale bar is 2 microns. (B) Frames from time-lapse showing gliding of contractomeres around an extruding cell (white asterisk) in a homeostatic mature monolayer. Contractomere-pairs flanking the apical junction between neighboring cells next to the extrusion event are marked by circles and squares with same color. During cell extrusion, one contractomere of the pair is stationary (circles) while the other contractomere (squares) move away to extend the junction. Merge image shows the absence of global junction movement. Lower left graph shows the distance travelled by the motile contractomeres of 3 contractomere pairs, one is immobile and the other is motile. Lower middle graph shows the distance between 2 immobile contractomeres in the same cells. Lower right graph shows the distance travelled all the motile contractomeres surrounding an extruding cell. The perimeter of the extruding cell is plotted to show the decrease in junctional length correlates with elongation of junctions between neighboring cells. Scale bar is 5 microns. (C) Graph summarizes the total net change in junctional lengths plotted in lower right graph of 5B. (D) Frames from time-lapse showing gliding of contractomeres around an extruding cell (asterisk) in a mature monolayer. During cell extrusion, contractomeres immediately surrounding the extruding cell (squares) move away from the immobile contractomeres (circles). Merge image shows the absence of global junction movement during cell extrusion. Scale bar is 5 microns.

These results support our hypothesis that contractomere is a motorized structure with actin motility function. Our observation is consistent with known motorized organelles with associated molecular motors myosin, kinesin, or dynein. The behavior of individual motorized organelle is dictated by its associated motor but not the behaviors of other motorized organelles in proximity. Thus, junction contractomeres are likely to behave independently of each other as autonomous motorized organelles.

### Blebbistatin blocks actin accumulation at the contractomere

The existence of contractomeres underscores a new way to think about junction processes in epithelial cell sheets. Myosin IIB, a vertebrate paralog of myosin II, has higher duty ratio and produces greater power than the ancestrally derived myosin IIA (Melli et al., 2018; Stam et al., 2015; Wang et al., 2003). Incorporating myosin IIB into a junctional complex would not only confer “motor” function but also support possessive movement on actin filaments. The goal for the remainder of this study is to determine whether contractomere is a new “organelle” that possess biochemical activities to “walk the junction”.

Previously, we reported that treating mature epithelial monolayers with latrunculin B, an actin monomer sequestration drug that promotes actin dissociation from filament ends, resulted in the disassembly of almost all actin structures except a latrunculin-resistant pool associated with alpha-actinin-4 at the apical junction (Tang and Brieher, 2012). We now realize that alpha-actinin-4-latrunculin-resistant structures are contractomeres (Fig.6A). We suspect that myosin II ATPase activity plays a role in actin accumulation since myosin IIB is enriched at contractomeres (Fig 1D & 6B). To develop this idea further, we assessed junctional actin using a combination of well-characterized inhibitors of actin dynamics. We found that blebbistatin, a myosin II ATPase inhibitor (Limouze et al., 2004; Ramamurthy et al., 2004), completely prevented the formation of latrunculin-resistant actin (Fig 6C), indicating a role of myosin II ATPase actin motility in this process.

**Figure 6.**
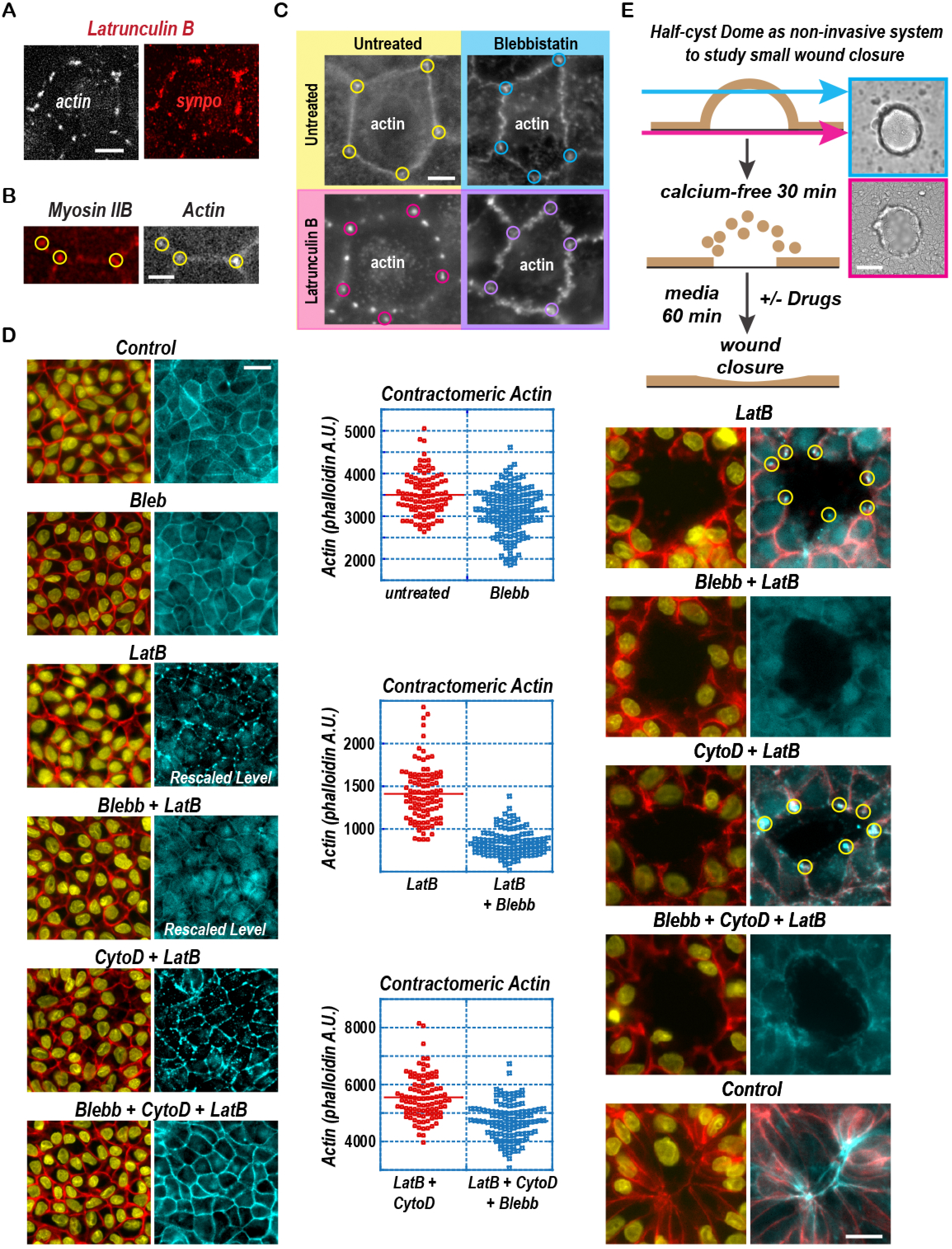
Contractomeric myosin II activity is linked to actin accumulation. (A) Immunofluorescence showing synaptopodin colocalizes with latrunculin-resistant actin at the apical junction. Scale bar is 2 microns. (B) Immunofluorescence showing myosin IIB colocalizes with actin puncta. Scale bar is 1 micron. (C) Myosin II inhibition prevents the formation of latrunculin (LatB)-resistant actin at the apical junction. Scale bar is 2 microns. (D) Myosin II inhibition prevents the formation of latrunculin (LatB)-resistant actin in the presence or absence of barbed-end capping by cytochalasin D (CytoD). Graphs show intensity measurement of actin on individual contractomeres. Bars mark the means. P<0.001 between untreated and blebbistatin treated groups for all 3 graphs. Scale bar is 10 microns. (E) Non-invasion wound healing assay showing the formation of latrunculin-resistant actin at wound edge, circled, which is inhibited by blebbistatin (Bleb). Latrunculin-resistant actin is formed after 1-hour incubation with cytochalasin D (CytoD) and latrunculin B (Lat B). In the absence of blebbistatin (Bleb), cytochalasin D (CytoD) or latrunculin B (Lat B), the wound is closed in an hour. Upper right panels show phase-contrast images of MDCK dome before wounding. Scale bars are 10 microns.

How does the contractomere accumulate actin in the presence of latrunculin B, which creates an environment that favors depolymerization of actin filaments? One possibility is that the contractomere can polymerize actin using latrunculin-bound actin. To eliminate this highly unlikely possibility, we blocked actin dynamics using cytochalasin D to prevent filament elongation from the barbed-ends (Fig 6D, bottom 2 rows). Strikingly, barbed-end capping by cytochalasin D enhanced actin accumulation at contractomeres under latrunculin B depolymerization environment. One interpretation of this result is that myosin IIB is capable of possessively “walk” on actin filament and “reel-in” actin filaments to form an actin ball at the contractomere. If this is true, blocking myosin II activity should prevent actin accumulation. Indeed, blebbistatin completely abolished latrunculin-induced actin accumulation at the contractomere, either in the presence or absence of cytochalasin D (Fig 6D, second, fourth, and sixth rows). These observations support our hypothesis that contractomeric myosin IIB can possessively “walk” on actin filaments, conferring actin motility activity to the contractomeric complex.

Actomyosin activity at ZO-1 apical junctions has previously been shown to involve in wound constriction (Tamada et al., 2007). To further explore the relationship between contractomere motility and junction constriction, we repeated the drug experiments on small wounds to determine the impact on wound closure (Fig 6E). Treatment of small wound with latrunculin B resulted in the formation of actin puncta around wound edges (Fig 6E, first and third rows). Actin puncta translocate within an hour to constrict the wound (Fig 6E, right panels), which is inhibited by blebbistatin (Fig 6E, second and fourth rows).

These drug studies raised an important question about the role of actin pointed-end at the contractomere. Latrunculin-resistant actin puncta are formed when the barbed-end were blocked by cytochalasin D, suggesting that the formation of latrunculin actin puncta is due to pointed-end depolymerization. These results implicate that the pointed-end of actin filaments is associated with the contractomere. Indeed, this argument is consistent with our previous findings showing that actin assembly at the contractomere requires Arp2/3, which binds to actin pointed-ends (Kannan and Tang, 2015). Collectively, these observations support our hypothesis that contractomeric myosin IIB acts as a barbed-end-directed motor to “reel-in” actin filaments that has barbed-ends facing out.

### Contractomere couples myosin activity and actin polymerization

The contractomere was originally characterized in a junction-enriched membrane fraction containing *de novo* actin assembly activity (Tang and Brieher, 2012). This activity requires Arp2/3-dependent actin nucleation and alpha-actinin-4-dependent actin polymerization. Using chemical crosslinking and alpha-actinin-4 as bait, we identified synaptopodin in the complex (Tang and Brieher, 2013). Subsequently, we showed that synaptopodin interacts with myosin II (Kannan and Tang, 2015), implicating a role of myosin II in contractomeric actin assembly *in vitro*. Here, we performed biochemical analysis to investigate the structure-function relationship between alpha-actinin-4, myosin IIB, and actin assembly using this purified membrane system that we had characterized extensively (Kannan and Tang, 2015; Tang and Brieher, 2012; Tang and Brieher, 2013). Briefly, actin assembly is initiated by the addition of 2 uM of fluorescently-labelled monomeric actin, a concentration substantially above the critical concentration of actin, to junction-enriched membranes in the presence of ATP.

To establish a role of myosin II in contractomeric actin assembly, we performed *in vitro* actin polymerization assays on junctional membranes that show colocalization of myosin IIB and synaptopodin with alpha-actinin-4 (Fig S9A). We found that contractomeric actin assembly is enriched at membrane sites marked by myosin IIB (Fig S9B). Assembly of actin on contractomeres *in vitro* was absent in the presence of blebbistatin, indicating an essential role of myosin II in this process (Fig S9C). Negative-staining electron microscopy (EM) of the membrane reaction revealed a ball of actin filaments rolled up in a ∼350-400 nm structure (Fig S9D, upper left panel). Since the “ball” of actin filaments obstructed the features of the contractomere, we lowered the among of actin in the reaction to just slightly above its critical concentration to limit actin filament formation. Under this condition, we observed small membrane organelles with multiple globular sites interacting with single actin filaments (Fig S9D, top right and lower panels).

A striking feature of the contractomere under EM is the lack of actin crosslinking. We only found contractomere interacting with single or parallel actin filament rather than a meshwork. Yet, actin assembly requires alpha-actinin-4, an actin crosslinking protein. Why wound actin assembly require an actin-crosslinking protein? Why do we not see actin networks at the contractomere? One possibility is that alpha-actinin-4 does not function as an actin crosslinker at the contractomere. To determine if crosslinking is necessary for contractomeric actin assembly, we performed a reconstitution assay by stripping the membranes with high salt to remove endogenous alpha-actinin-4 and replacing the reaction with recombinant alpha-actinin-4, an assay we had previously developed for the identification of alpha-acitnin-4 and synaptopodin (Tang and Brieher, 2012; Tang and Brieher, 2013). Using the reconstitution assay, we tested different truncations of alpha-actinin-4 to determine whether actin crosslinking is necessary for actin assembly.

Alpha-actinin exists as an anti-parallel dimer ∼36 nm in length (Meyer and Aebi, 1990) containing 2 actin-binding calponin-homology domains flanked by 4 spectrin-repeats that support high affinity anti-parallel dimer formation (Liu et al., 2004). We generated recombinant alpha-actinin-4 that has one actin-binding domain (ABD) instead of two actin-binding domains, thus cannot crosslink actin filaments into networks. For comparison, we generated alpha-actinin-4 that has no actin-binding domain or missing the spectrin repeats (Fig 7A). After we obtained recombinant alpha-actinin-4 proteins, we labelled them with a fluorophore so that we can track their targeting to the contractomere (Fig 7B). We found that alpha-actinin-4 recruitment to the contractomere does not require actin-binding; alpha-actinin-4 missing both actin-binding domains can still target (Fig 7B). This is in sharp contrast to other actomyosin structures such as stress fiber or actin meshwork where binding to actin filaments is necessary for their assembly. Moreover, alpha-actinin-4 with only one actin-binding domain would support contractomeric actin assembly to the same extent as alpha-actinin-4 with two actin-binding domains (Fig 7C). By contrast, spectrin repeats or actin-binding domain alone cannot support actin assembly, indicating that both contractomere targeting and actin-binding are necessary (Fig 7B-C). These results demonstrated that contractomeres exhibit novel and unique biochemistry distinct from other known alpha-actinin structures.

**Figure 7.**
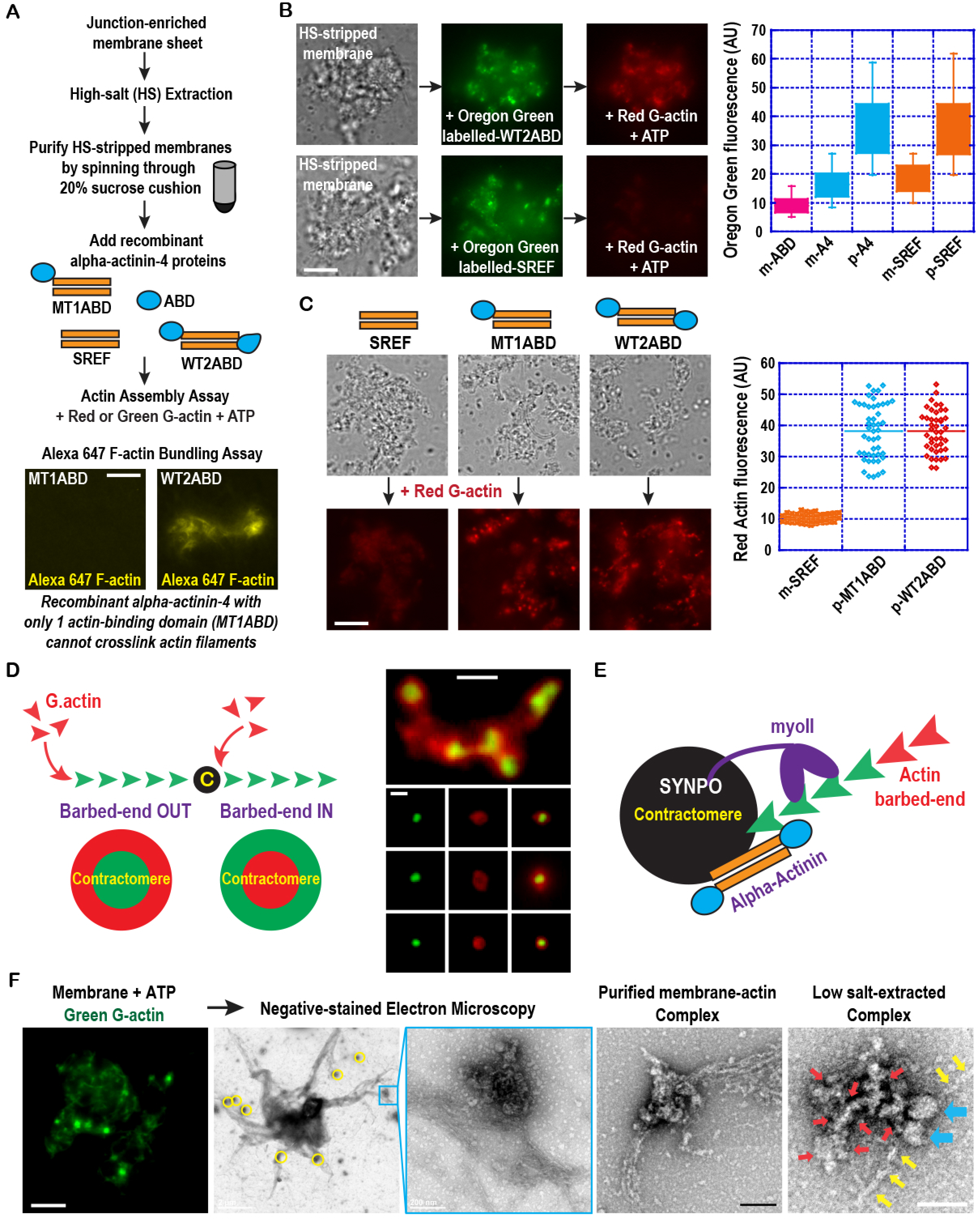
Contractomere alpha-actinin-4 associates with actin filaments near the pointed. (A) Reconstitution actin assembly assay using stripped membranes and recombinant alpha-actinin-4 full-length or truncated proteins. Bottom panels shows that alpha-actinin-4 with one actin-binding domain failed to crosslink Alexa 647-labelled actin filaments into bundles that can be readily visualized under the light microscope. Scale bar is 2 microns. (B) Actin assembly on contractomere requires alpha-actinin-4 binding to actin. Reconstitution assay showing alpha-actinin-4 lacking the actin-binding domains was recruited to the contractomere but failed to support actin assembly. Graph shows intensity measurement of Oregon green-labelled alpha-actinin-4 on individual contractomeres; boxes represent 75 percentile and error bars are standard deviation. Membrane-bound signals are m-ABD, m-A4, m-SREF and contractomere signals are p-A4 and p-SREF. P < 0.001 between m-A4 and p-A4 and between m-SREF and p-SREF. P>0.01 between p-A4 and p-SREF. Scale bar is 2 microns. (C) Contractomeric actin assembly requires one actin-binding domain of alpha-actinin-4. Actin assembly using alpha-actinin-4 with either one or two actin domains is the same. Graph shows intensity measurement of rhodamine-labelled actin on individual contractomeres. Bars mark the means. p-MT1ABD and p-MT2ABD refer to contractomeric signals. P< 0.001 between p-MT1ABD and p-SREF. P< 0.001 between p-MT2ABD and p-SREF. P>0.01 between p-MT1ABD and p-MT2ABD. Scale bar is 2 microns. (D) Pulse-chase actin assembly using Oregon green-labelled G-actin followed by rhodamine-labelled G-actin (see Results & Methods). Barbed-end addition of red G-actin shows that actin-barbed ends are facing away from the contractomere. Scale bars are 500 nm. (E) Summary of result and working model. Contractomeric alpha-actinin-4 can hold onto the side of an actin filament from the pointed-end which is facing the contractomere. Myosin IIB can walk towards the barbed-end of the actin filament. (E) Negative-stain electron microscopy showing actin filaments associated with contractomeres and a sub-complex of contractomere containing a myosin II monomer (see Methods). An actin filament (yellow arrows) is interacting with both motor heads of the myosin II monomer (blue arrows). The tail region of the myosin II monomer interacts with electron-dense materials (red arrows), analogous to “stuff” associated with tails of myosin II monomers seen in cells (see Results & Discussion). Scale bars in 2 left panels are 2 microns. Scale bars in middle and middle-right 2 panels are 200 nm. Scale bar on right panel is 50 nm.

In the above drug studies in cells, we have mentioned the possibility that actin pointed-end is facing towards the contractomere whereas filament barbed-end needs to face away for the barbed-end-directed myosin IIB motor to “reel-in” actin (Fig 6D-E). To test whether actin barbed-end is facing away, we used a 2-color “pulse-chase” assay to determine the location of actin addition (Fig 7D). If actin adds next to the contractomere, the barbed end is likely to localize on the contractomere. However, if actin adds to the outside of polymerizing actin, the barbed-end is likely to face outwards. We initiated the actin assembly reaction on contractomeres using a low concentration of FITC-labelled G-actin to decrease the rate and extent of actin assembly. After removing unincorporated FITC-labelled G-actin, we continue the reaction using TRITC-labelled G-actin (Fig 7D). We imaged the reaction using wide-field microcopy and found that TRITC-labelled actin is incorporated on the outside of FITC-labelled filaments, consistent with our hypothesis that barbed-ends are facing outwards (Fig 7D, right panels). These observations support a previous observation that actin pointed-ends are pointing towards the apical junction in epithelial cells (Madara, 1987).

Calponin homology domains are side-binding domains, thus alpha-actinin-4 is likely to hold onto the side of an actin filament close to the pointed-end at the contractomere (Fig 7E). In this configuration, contractomeric myosin IIB can walk towards the barbed-ends of the actin filaments, pulling filaments towards the contractomere. Since latrunculin can depolymerize actin filaments from pointed-ends and barbed-ends, it can disrupt pointed-end interactions with alpha-actinin-4 as well as barbed-ends interactions with junctional proteins on the junction. When these interactions are destabilized by latrunculin, myosin IIB activities can “reel-in” actin in an unregulated manner, rolling the filaments into a ball. The degree of actin ball formation would depend on the speed of the myosin IIB motor and the rate of barbed-end depolymerization. At the apical junction, actin barbed-ends are capped by CD2AP (Tang and Brieher, 2013). Capping actin barbed-ends would block actin depolymerization, allowing the filaments to be collected at the contractomere by myosin IIB. This is consistent with previous finding that knockdown of CD2AP prevented the formation of latrunculin-resistant actin (Tang and Brieher, 2013). Collectively, our results showed that contractomeric alpha-actinin-4 holds onto actin filaments at their pointed-ends while the barbed-ends are stabilized by CD2AP during latrunculin treatment. Our hypothesis is consistent with previous observation showing functional interactions between synaptopodin and CD2AP (Huber et al., 2006).

One unsettling feature of contractomere under EM is the lack of myosin II minifilaments (Fig S9D). A myosin II minifilament is ∼300 nm in length characterized by a dumbbell-shape with myosin heads spraying out at both ends and a bare mid-zone (Liu et al., 2017; Liu et al., 2018). Yet, we could not find a structure resembling a myosin II minifilament despite screening 10 membrane preparations with contractomeric actin assembly activities. Purified contractomere is less than 250 nm in dimension (Fig S9D, upper right and lower left panels) (Kannan and Tang, 2015), which is too small to hold a myosin II minifilament. In fact, extraction of junctional membranes with non-ionic detergent revealed a contractomere no bigger than 150 nm in dimension but still can interact with actin filaments at multiple sites (Fig S9D, lower right panel) (Kannan and Tang, 2015).

Recently, monomeric myosin II has emerged as an important player in diverse cellular functions including focal adhesion initiation, cell migration, Golgi dynamics, and exocytosis (Aoki et al., 2010; Ganguly et al., 1992; Kiboku et al., 2013; Shutova et al., 2017; Shutova et al., 2014). Here, we used a modified protocol to assess whether contractomere might contain monomeric myosin II (Fig 7F). We empirically tested extraction conditions by varying the concentrations of chaotropic agents and screened for conditions that may break the contractomere into subcomplexes. Using a combination of mechanical shearing and extractions with low salt and basic pH buffers; we found a subcomplex that contains a structure resembling a myosin monomer (Fig. 7F, right panel, yellow arrows). This is an interesting finding because the pKa of synaptopodin is >10, indicating that basic pH might be a condition to break up its interaction with contractomeric proteins. The subcomplex appears to show a myosin monomer bound to electron dense materials at its coiled-coil tail and interacting with an actin filament at the head domains (Fig. 7F, right panel, red arrows). These observations are consistent with recently published works by the Svitkina group showing electron-dense “stuff” bound to the coiled-coil tail of monomeric myosin II in cells (Shutova et al., 2017; Shutova et al., 2014).

Based on our hypothesis, myosin IIB barbed-end-directed motility would shove actin filaments into the contractomere. We predict that shoving actin at alpha-actinin-4 located closer to the pointed-end of actin filaments would exert force on alpha-actinin-4 within the contractomere (Fig 7E). To test this hypothesis, we have designed new FRET-based alpha-actinin-4 sensors to measure changes in force at the contractomere.

### Alpha-actinin-4 experiences force at the contractomere

We had previously used a FRET-based force sensor, sstFRET, to measure E-cadherin and myosin-1c tension in epithelial cell sheets, indicating that the sensor is capable of registering cellular forces (Kannan and Tang, 2018). The sensor contains a single spectrin repeat flanked by a venus(VFP)-cerulean(CFP) FRET-pair, which has been calibrated to report on ∼6 pN of force with a 50% change in FRET (Meng and Sachs, 2011). We inserted the sstFRET cassette away from the actin-binding regions of alpha-actinin-4 to avoid interfering with actin binding but at a position where we think might report on changes in force when alpha-actinin-4 is bound to actin filaments (Fig. 9A). The spectrin-repeats of alpha-actinin can interact with many junctional proteins including alpha-catenin, vinculin, synaptopodin, and ArgBP2 (Asanuma et al., 2005; Bois et al., 2005; Chen et al., 2006; Knudsen et al., 1995). To avoid interfering with these interactions, we positioned the sstFRET cassette within a flexible linker region between the 2nd and 3rd spectrin repeats at residue 522 (alpha-actinin-4-sstFRET522) such that the anti-parallel interactions and the dimer interfaces formed by the spectrin-repeats are preserved (Fig 8B). We confirmed the expression of alpha-actinin-4-sstFRET522 by western blots and showed that it does not interact with endogenous alpha-actinin-4 by immunoprecipitation (Fig 8B).

**Figure 8.**
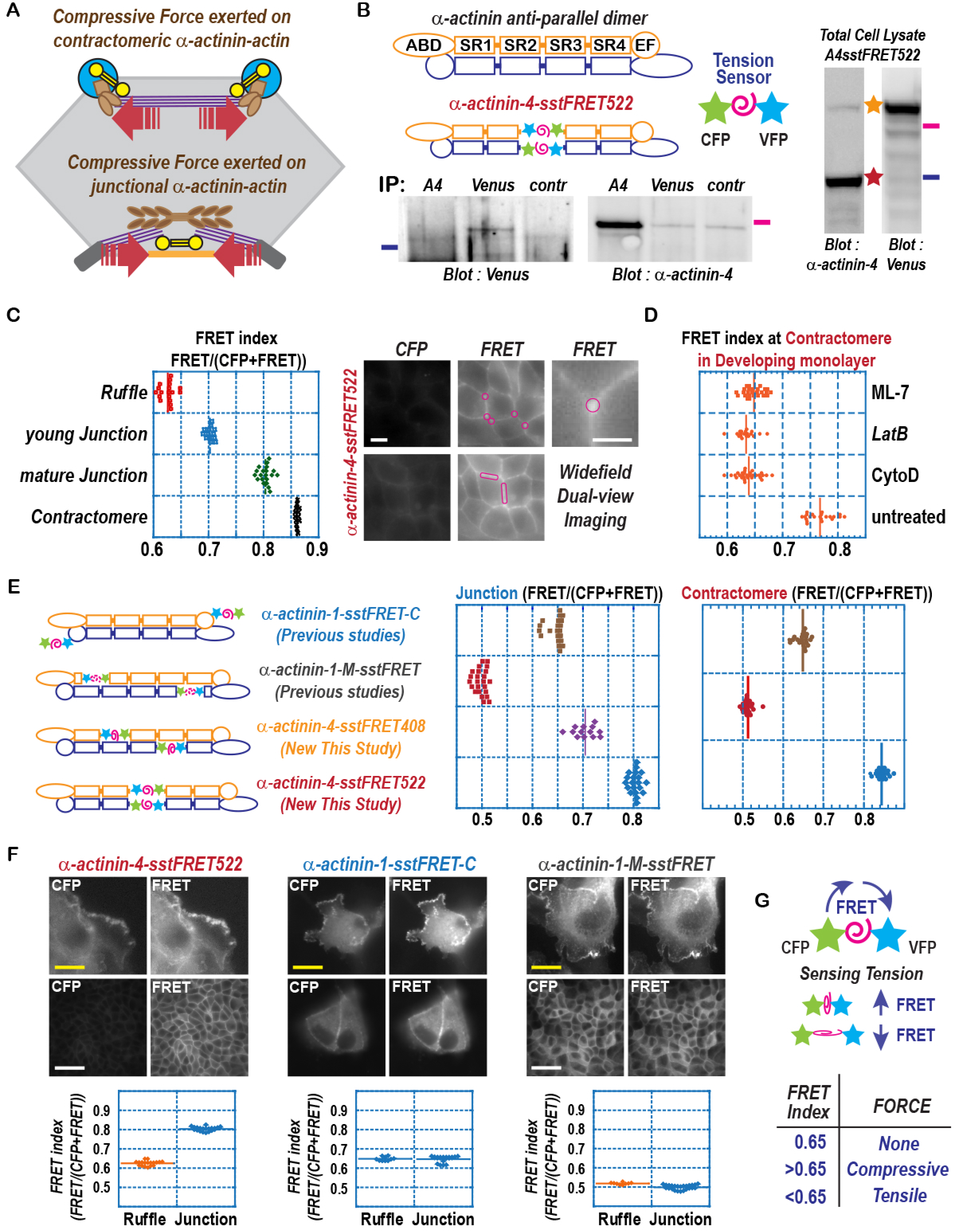
Contractomeric alpha-actinin-4 experiences actomyosin force. (A) A working hypothesis showing contractomeric alpha-actinin might experience actomyosin-dependent force. (B) Design and expression of an alpha-actinin-4 force sensor (alpha-actinin-4-sstFRET522). Right panels are western blots for venus and alpha-actinin-4 showing expression of alpha-actinin-4-sstFRET522 in MDCK cells (see Methods). Lower panels are western blots of an immunoprecipitation showing negligible interaction between alpha-actinin-4-sstFRET522 and endogenous alpha-actinin-4. Markers are 135 kD in green and 100 kD in orange. (C) FRET measurements (see Methods) showing low FRET at protrusions and high FRET at contractomeres. Right panels show the areas for contractomere and junction measurements. Scale bar is 10 microns. (D) Inhibition of actomyosin reduces alpha-actinin-4-sstFRET522 FRET index at the contractomere. P<0.001 between untreated and drug treated groups. (E) Comparison of 4 alpha-actinin FRET-based sensors containing the force-sensing cassette, sstFRET, at different position of alpha-actinin. When the force sensor is located at the C-terminus (alpha-actinin-1-sstFRET-C), no force can be applied to the sensor and the FRET index is 0.65 reflecting baseline FRET in cells. Alpha-actinin-1-M-sstFRET reported a tensile force at the junction whereas alpha-actinin-4-sstFRET408 and alpha-actinin-4-sstFRET522 reported compressive forces. (F) FRET difference between protrusive cell edge and contractile cell junction using alpha-actinin-4-sstFRET522 but not alpha-actinin-1-sstFRET-C or alpha-actinin-1-M-sstFRET. P<0.001 between ruffle and junction measurements. Scale bars are 5 microns. (G) FRET readings as proxies for tensile and compressive forces.

**Figure 9.**
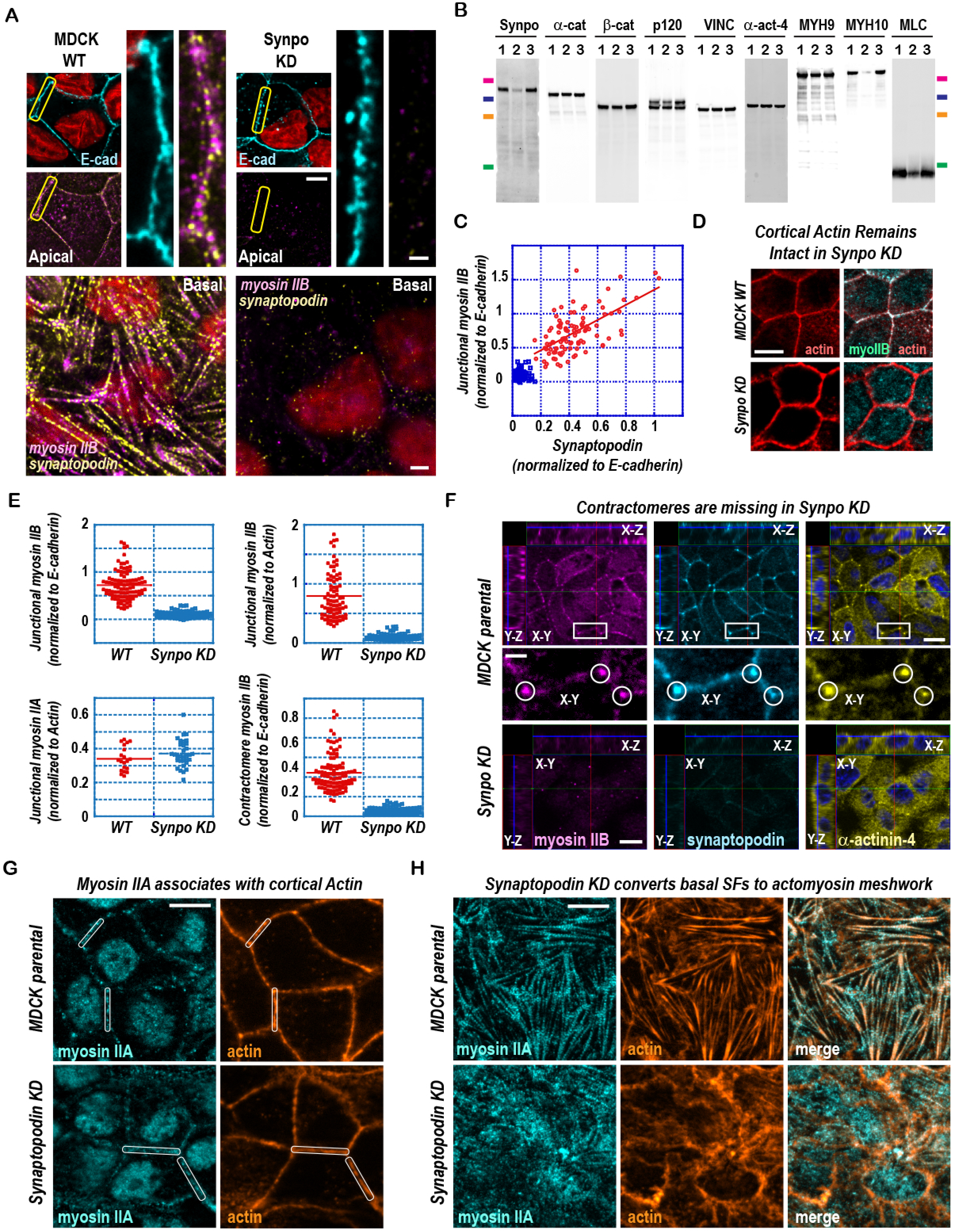
Contractomeres and stress fibers are missing in synaptopodin knockdown cells. (A) Immunofluorescence of MDCK wild type (WT) and synaptopodin knockdown (Synpo KD) cells. Apical and basal stress fibers are missing in synaptopodin knockdown cells. Scale bars are 1 micron. (B) Western blots showing reduced cellular levels of myosin IIB and myosin light chain in synaptopodin knockdown cells. Lanes 1 and 3 are whole cell lysate of MDCK parental cells. Lane 2 is whole cell lysate of synaptopodin knockdown cells. Markers are 150, 100, 75, 25 kD. (C) Scatter plot showing relationship between synaptopodin and myosin IIB levels in MDCK WT (red dots) and Synpo KD (blue squares) at the apical junction. Pearson correlation coefficient is 0.63 between synaptopodin and myosin IIB in WT junction. (D) Myosin IIB is absent from the apical junction in synaptopodin knockdown cells despite the presence of junctional actin. Scale bar is 5 microns. (E) Synaptopodin knockdown (Synpo KD) specifically reduced myosin IIB but not myosin IIA. Measurement of junctional and contractomeric intensities of myosin IIA and IIB immunofluorescence. P<0.001 between WT and Synpo KD for myosin IIB. P>0.01 between WT and Synpo KD for myosin IIA. (F) Contractomeres at apical junctional vertices (circled) are missing in synaptopodin knockdown cells. Z-stacks are shown in X-Y, Y-Z, and X-Y axes. In synaptopodin knockdown cells, myosin IIB level was lower and alpha-actinin-4 became cytoplasmic. Scale bars are 5 microns. (G) Myosin IIA is present at the apical junction with actin in synaptopodin knockdown monolayers. Scale bar is 5 microns. (H) Basal myosin IIA stress fibers are converted to into myosin IIA meshworks in synaptopodin knockdown cells. Scale bar is 5 microns.

Using a dual-view beam splitter to measure CFP and FRET simultaneously, we compared alpha-actinin-4-sstFRET522 FRET at different cellular locations including cell protrusions, cytoplasm, linear junction, and contractomere (Fig. 8C). We found that alpha-actinin-4-sstFRET522 FRET was substantially higher when measured at the contractomere than at membrane ruffle, a protrusive structure consisted of actin meshworks. When we inhibited myosin II or actin dynamics with myosin light chain inhibitor ML-7, latrunculin B or cytochalasin, the FRET signal at the contractomere is drastically reduced (Fig 8D). These observations suggest that myosin II at the contractomere is capable of exerting force on actin filaments, which is measurable by our new alpha-actinin-4-sstFRET522 sensor.

In addition to alpha-actinin-4-sstFRET522, we compared 3 other sstFRET-based sensors (Fig 8E-G). One sensor has sstFRET inserted at the C-terminus (alpha-actinin-1-sstFRET-C), thus cannot experience externally applied force. Measurement of alpha-actinin-1-sstFRET-C showed a FRET index of ∼0.65, representing the background FRET in cytoplasm. FRET index of ∼0.65 is comparable to alpha-actinin-4-sstFRET522 FRET when actomyosin dynamics were inhibited, indicating a lack of force under these conditions (Fig 8C, graph). FRET index of ∼0.65 is also comparable to alpha-actinin-4-sstFRET522 FRET measured at membrane ruffles, indicating that alpha-actinin-4 does not experience measurable force at protrusion either. Two additional sensors we tried had sstFRET inserted within the first spectrin repeat of alpha-actinin-1 (alpha-actinin-1-M-sstFRET) or between the 1st and 2nd spectrin repeats of alpha-actinin-4 (alpha-actinin-4-sstFRET408). One of the sensors reported an increase in FRET and the other a decrease in FRET when compared to alpha-actinin-1-sstFRET-C, indicating that the sstFRET FRET-pair might be stretched or compressed when inserted at these positions in alpha-actinin (Fig 8E). These sensors also failed to report a change in FRET between protrusive and contractile structures (Fig 8F-G), indicating that they are not able to measure force.

### Knockdown of synaptopodin abolishes stress fibers and contractomeres

We have identified stress fiber and contractomere as synaptopodin structures at the apical junction. To assess whether these structures require synaptopodin for assembly, we knocked down synaptopodin in MDCK cells. In developing monolayer, stress fibers were absent in synaptopodin knockdown cells (Fig. 9A). Moreover, synaptopodin knockdown selectively reduced myosin IIB and myosin light chain levels without affecting junctional components including alpha-catenin, beta-catenin, p120-catenin, vinculin, alpha-actinin-4, or myosin IIA (Fig 9B-E). In mature synaptopodin knockdown cells, contractomeres were missing and alpha-actinin-4 became cytoplasmic (Fig 9F). In contrast, myosin IIA and actin at the junctional cortex were not affected (Fig 9G). However, stress fibers on the basal cell surface are now converted into a myosin IIA-actomyosin meshwork (Fig 9H).

Synaptopodin is essential for the maintenance of permeability barrier in multiple systems including MDCK cells, T84 intestinal cells, mouse intestine, and kidney podocytes (Kannan and Tang, 2015; Ning et al., 2020; Ning et al., 2021; Wang et al., 2020). However, the mechanism for its protective function is not known. Now we know that synaptopodin depleted cells are missing stress fibers and contractomeres. Synaptopodin depletion decreases the rate of wound migration in endothelial cells, epithelial cells, and podocytes (Mun et al., 2014; Ning et al., 2020; Ning et al., 2021; Rochman et al., 2017), which is consistent with the role of stress fibers in lamella formation in migrating cells (Burridge and Guilluy, 2016). Several studies have also shown that epithelial integrity is disrupted to a greater extent in synaptopodin depleted cells when challenged by chemical insult, indicating a protective role and a survival advantage provided by synaptopodin. These advantages could be due to the ability of contractomeres to quickly close wounds and preserve the epithelial permeability barrier during cell extrusion.

Synaptopodin knockout mice resulted in down-regulation of RhoA in adult animals (Asanuma et al., 2006; Ning et al., 2020). RhoA is a regulator of stress fiber assembly and contractility (Ridley and Hall, 1992), thus could play a role in synaptopodin function. However, in MDCK cells, synaptopodin knockdown did not affect the level or activity of RhoA (Fig S10A-B). Nevertheless, phospho-myosin light chain is greatly compromised by synaptopodin depletion (Fig S9B & 10B), indicating that synaptopodin in MDCK cells might use an alternative pathway to control actomyosin structures and contractility. Sequence analysis of synaptopodin indicates that synaptopodin contains putative binding sites for ArgBP2/SORB2 (Scansite, MIT), a vertebrate paralog of the conserved SORBS1/ponsin (Ichikawa et al., 2017). ArgBP2/SORB2 regulates stress fiber contractility whereas SORBS1/ponsin regulates focal adhesion stability (Anekal et al., 2015; Ichikawa et al., 2017). ArgBP2/SORB2 targets to the apical junction and stress fiber in an alpha-actinin-dependent fashion and has previously been shown to interact directly with alpha-actinin-4 (Anekal et al., 2015; Fredriksson-Lidman et al., 2017). ArgBP2 also interacts with WAVE2 (Cestra et al., 2005), which regulate actin dynamics at the epithelial junction via Arp2/3 (Nakanishi et al., 2007; Verma et al., 2012). Here, we show that depletion of synaptopodin significantly reduced the level of ArgBP2/SORB2 (Fig S10C), implicating a role of synaptopodin/ArgBP2 in actomyosin regulation in a RhoA-independent manner.

Collectively, our results underscore the roles of vertebrate-specific synaptopodin not only in the assembly of vertebrate-specific actomyosin structures, stress fibers and contractomeres, but also in the regulation of vertebrate-specific ArgBP2/SORBS2 mechanotransduction pathway.

## Discussion

In this study, we have described a 2 novel synaptopodin-dependent actomyosin structures: apical stress fiber and contractomere (Fig 10). The first structure, which we have named synaptopodin apical stress fiber, is consisted of alternating and periodic arrangements of myosin II, synaptopodin, alpha-actinin, and actin. Unlike basal stress fibers that are inserted at vinculin-enriched cell-cell adhesions, apical stress fibers are inserted at E-cadherin & ZO-1-decorated apical junctions. When an apical stress fiber is attached head-on to the junction, orthogonal force can apply to the junctional complex. By contrast, if the apical stress fiber is attached side-on via synaptopodin linkers, the linear junction would experience parallel force. Contractions of parallel stress fibers are associated with lateral movement and clustering of junctional complexes whereas contraction of orthogonal stress fibers shortens the junction.

**Figure 10.**
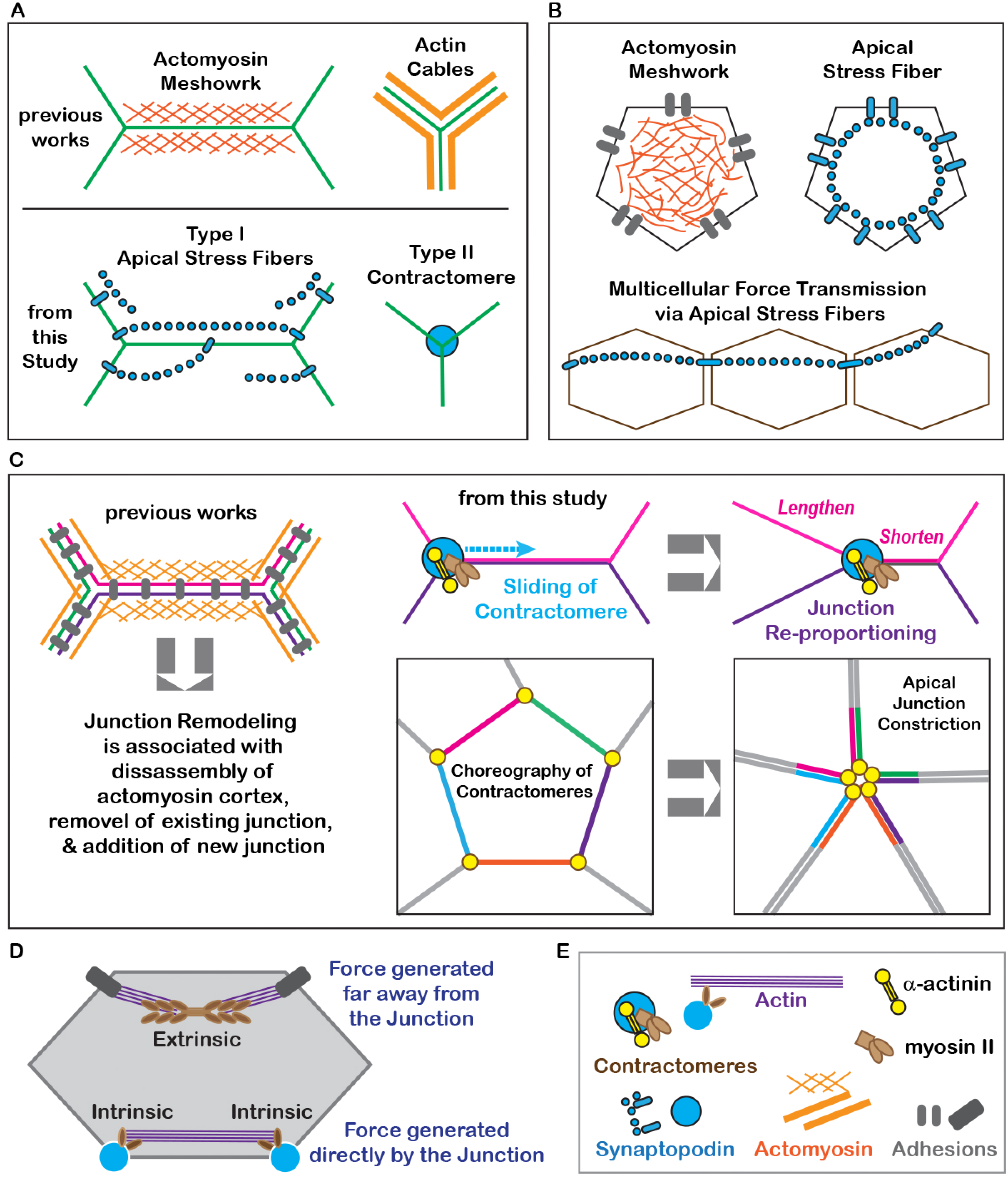
Actomyosin structures at the apical junction. (A) Comparison of actomyosin meshwork and cables with apical stress fiber and contractomere. Apical stress fibers can selectively link neighboring junctions as well as junctions from opposite sides of a cell. Contractomere is a unique actomyosin structure that contains non-filamentous myosin II. (B) Comparison between actomyosin meshwork and apical stress fibers is shown at the top panels. Apical stress fibers inserted at cell-cell adhesion can propagate force and generate tissue-level tension over many cells in an epithelial monolayer, the bottom panel. (C) Shortening and lengthening the junction by “walking” the contractomere. Motility of contractomeres contributes to apical junction constriction during cell extrusion and pure-string wound closure. The prevailing model of junction remodeling requires disassembly of the actomyosin cortex and endocytosis of existing junction to shorten a junction. (D) Contractomere is directly powered by myosin II and actin polymerization to generate force locally at the junction while apical stress fiber powered by myosin II are at a distance away from the junction. Contractomeres generate intrinsic force whereas apical stress fibers or actomyosin meshworks generate extrinsic force with respect to the junction. (E) Key to cartoon.

The second structure is a novel macromolecular assembly ∼150-200 nm in dimension containing myosin II, synaptopodin, and a-actinin-4. We named this structure “contractomere” for its biochemical and structural properties; “contracto” refers to the fact that it uses myosin II motor activity for actin motility function and “mere” refers to it being a minimal macromolecular complex to support its functions. We demonstrated that contractomere uses a novel mechanism by harnessing the energy from actin and myosin ATPases to support junction dynamics.

Contractomere exists as a standalone structure even after the removal of endogenous actin and can be isolated from tissues or cultured epithelial cells. This is in contrast to stress fibers or actomyosin networks where the interaction between actin filament and myosin II minifilament is the molecular basis for assembly and necessary for their structural integrity. Contractomere can glide along the junction to zip-up the plasma membranes of neighboring cells, creating new cell-cell interface. Motility of contractomeres during cell extrusion allow junction constriction while maintaining epithelial barrier integrity, thus provide a protective function in homeostatic epithelia.

We propose a model by which myosin II within the contractomere can “walk” on actin filaments and pull actin towards the center of the contractomere. Since myosin II is a barbed-end-directed motor, our model requires actin barbed-ends to face away from the contractomere for the motor to slide actin filaments in the direction of the contractomere. Our finding is opposite of what we would expect because myosin II contractility has always been thought to pull on the junction from a distant location.

To investigate our hypothesis further, we generated a novel alpha-actinin-4 FRET tension sensor to monitor forces at the contractomere. We showed that alpha-actinin-4 experiences changes in force in an actomyosin-dependent manner. These observations are consistent with myosin II pulling on actin filaments, shoving actin filaments toward the contractomere and exerting a compressive force on aalphaactinin-4 located within the contractomere.

Lastly, we showed that the formation of these 2 actomyosin structures require a vertebrate-specific protein called synaptopodin. Depletion of synaptopodin prevented the assembly of apical stress fiber and contractomere. Importantly, synaptopodin depletion resulted in the conversion of stress fiber into a meshwork-type actomyosin organization. Our results not only revealed that synaptopodin is a regulator of stress fiber assembly but that the type of actomyosin assemblies inside the cell can be tuned by synaptopodin.

### Re-defining actomyosin structures of the epithelial junction

The prevailing models of actomyosin structures at the epithelial junction describe 3 different actomyosin organizations for force generation, all of them depend on bipolar myosin II minifilaments (Fig 10A-B). The first model describes a sub-membrane cortical actomyosin II network (Fig 10A, upper left). The second model describes parallel actin cables bundled by actomyosin II (Fig 10A, upper right). The third model describes an isotropic actomyosin II meshwork on the apical cortex (Fig 10B, upper left). All 3 structures are likely to exert orthogonal force on the junction. However, they fall short in explaining how the junction can shrink during wound constriction, cell extrusion, or intercellular rearrangements. The 2 major issues are as follows.

The first issue is the fluid mosaic problem. In the first and second models stated above, force is exerted parallel to the junction, pulling adhesion complexes on the plane of the membrane. Parallel force would move adhesion complexes on the plane of the plasma membrane and potentially causing them to cluster. This mechanism will leave the lipids of the plasma membrane untouched rather than shrinking the junction. In the third model stated above, contraction of apical isotropic meshwork would pull on the junction perpendicularly, producing net zero parallel force which have no effect on junctional length.

The second issue is the 300 nm problem. Bipolar myosin II minifilaments are >300 nm long and geometrically cannot close the junction. If each linear junction has only one myosin II minifilament, a cell with 5 sides will be stuck with a hole >500 nm in diameter at the end of junction constriction. In Drosophila, apical constriction is arrested with a hole during wound healing, which must be closed by protrusion and cell migration (Abreu-Blanco et al., 2012; Wood et al., 2002). However, contractomere does not have the 300 nm geometric constrain of the myosin II minifilament and can constrict the junction completely without leaving a hole (Fig 10C, lower right panels).

A new concept emerged from this study is the origin of force. Force production by the contractomere is an intrinsic process since actin assembly and myosin II ATPase occur directly at the junctional complex (Fig 10D). By contrast, apical stress fibers or cortical actin meshworks are cytoplasmic structures. Force generated by these structures are extrinsic to the junctional complex and must be relayed via other protein assemblies. The existence of 2 independent force-production regimes increases the complexity of vertebrate mechanoregulation and the ability for epithelial cells to differentially regulate junction processes.

### Regulation of actomyosin structures at the epithelial junction

Apical stress fiber, contractomere, and actomyosin meshwork are 3 different force-generating structures, contributing to distinct force-dependent processes at the junction. The 3 regimes allow independent regulations and force transmission pathways that are structurally and functionally segregated. Previous studies had shown that myosin IIA structures are primarily controlled by Rho/Rock/MLC whereas myosin IIB structures are mainly regulated by Ca^++^/calmodulin/MLCK (Chang and Kumar, 2015; Kassianidou et al., 2017; Kuragano et al., 2018; Morin et al., 2014; Totsukawa et al., 2004). Phosphorylation studies using purified myosin II indicate that MLCK phosphorylates myosin IIB more efficiently than ROCK, whereas the reverse is true for myosin IIA (Amano et al., 1996; Sandquist et al., 2006). Furthermore, MLCK is >50 times more efficient in phosphorylating MLC than ROCK (Bresnick, 1999). Here, we show that ArgBP2/SORB2 may be another player in regulating actomyosin structures. A future goal is to determine the contribution of these pathways in the regulation and assembly of apical stress fiber and contractomere.

Myosin II bipolar minifilament has a characteristic dumbbell structure ∼300 nm in length consisted of a bare mid-zone and myosin heads spraying out from 2 ends. In purified junctional membranes, myosin II filamentous structure had not been observed. Instead, we found a compact macromolecular complex ∼150-250 nm in dimension (Kannan and Tang, 2015). Dissociation of the contractomeric complex using mechanical and biochemically perturbations showed that the coiled-coil tail of myosin II monomer was bound to proteins. These findings are consistent with previous reports by the Svitkina group describing the existence of electron-dense material surrounding the coiled-coil region of unfolded monomeric myosin II (Shutova et al., 2017; Shutova et al., 2014). The coiled-coil region has also been shown to play a role in targeting myosin IIB to secretory granules independent of actin (Milberg et al., 2017). These previous observations and our current results add to the growing number of studies showing that myosin IIB localization to different structures does not require actin binding (Badirou et al., 2014; Beach and Egelhoff, 2009; Fanning et al., 2012; Liu et al., 2016; Rosenberg et al., 2008; Roy et al., 2016). Thus, the interaction between synaptopodin and myosin IIB could potentially recruit myosin IIB to the contractomere. One of our future goals is to determine whether synaptopodin bind to the coiled-coil region of myosin IIB and whether the interaction is responsible for myosin IIB recruitment to the contractomere.

The Svitkina group has demonstrated that myosin II monomers are phosphorylated inside cells (Shutova et al., 2017; Shutova et al., 2014). Myosin II phosphorylation activates myosin II, converting myosin II from an inactive autoinhibited state into a constitutively active motor. Consistent with their observations, the Korn group has shown that recombinant myosin II monomers that are in the unfolded state can be phosphorylated *in vitro* (Liu et al., 2017; Liu et al., 2016; Liu et al., 2018). Collectively, these studies show that the motor function of unfolded monomers can be controlled by phosphorylation similar to filamentous myosin II. Importantly, phosphorylated unfolded monomeric myosin II are likely to be constitutively active with actin motility activity. Multiple phosphorylation sites have been identified on the coiled-coil region of myosin IIB, and phosphorylation of these sites by PKC-zeta and CK-2 promotes disassembly of myosin II minifilaments and formation of myosin II monomers (Dulyaninova and Bresnick, 2013; Even-Faitelson and Ravid, 2006; Juanes-Garcia et al., 2015; Murakami et al., 1998; Murakami et al., 1984; Vicente-Manzanares et al., 2009). PKC-zeta and CK-2 are kinases that regulate the development, stability, and maintenance of epithelial junctions (Dorfel et al., 2013; Eckert et al., 2005; Helfrich et al., 2007; Raleigh et al., 2011). Proteomics analysis of PKC-zeta membrane complexes have previously identified synaptopodin (Tang, 2006). Thus, regulation of myosin IIB by PKC-zeta and CK-2 might contribute to junction maturation by converting myosin II filaments into monomers at the apical junction. Our results show that apical stress fiber, which is formed by bipolar myosin II filaments, dissolves during junction maturation with concomitant formation of contractomere, formed by monomeric myosin IIB. The dissolution of apical stress fibers is specific at the apical junction because basal stress fibers remain intact, indicating that the apical junction contains regulators for the conversion process. Local regulation at the apical junction would provide an immediate pool of monomeric myosin IIB for the assembly of contractomeres. This is in contrast to the current paradigm that monomeric myosin II is the inactive reserved pool for myosin II minifilament assembly. One of our future goals is to determine the mechanism responsible for converting bipolar filaments into monomeric myosin IIB at the apical junction.

## Materials and Methods

### Antibodies and reagents

Primary antibodies were purchased from commercial sources and custom-generated. Rabbit and goat polyclonal antibodies to synaptopodin were raised against synthetic peptides corresponding to 4 regions of human synaptopodin X1 (NCBI Reference Sequence: XP_016864497.1): aa164-180 (PSSNSRGVQLFNRRRQR), aa675-684 (QQESAPRDRA), aa899-918 (SPRAKQAPRPSFSTRNAGIE), and aa1123-1143 (CPRGWNGSLRLKRGSLPAEAS). The peptides were synthesized and coupled to KLH before injection into the animals (Pacific immunology). Serum reactivity to the peptides were assessed by ELISA (Pacific immunology). Rabbit polyclonal antibodies to α-actinin-4 were raised in house against a synthetic peptide corresponding to aa 8-24 of human α-actinin-4, NQSYQYGPSSAGNGAGC, that has been coupled to KLH. Antibodies to α-actinin-4 (sc-3933495, mouse), beta-catenin (sc-7963, mouse), synaptopodin (sc-515842, mouse; sc-21537, goat), α-catenin (sc-9988, mouse), p120 (sc-13957, rabbit), vinculin (sc-5573, rabbit), myosin IIA (sc-47201, rabbit), myosin IIB (sc-376942, mouse), RhoA (sc-418), MRIP (sc-135494) and ArgBP2 (sc-514671) were purchased from Santa Cruz Biotechnology. Rabbit polyclonal antibodies to myosin IIA (19098) and myosin IIB (19099) were purchased from Biolegend. Antibodies to phospho-myosin light chain 2 (Thr18/Ser19) were purchased from Cell Signaling Technology (#3674, rabbit). Secondary antibodies were purchased from Bio-Rad Laboratories (HRP goat anti–rabbit), Santa Cruz Biotechnology (HRP rabbit anti–goat and goat-anti-mouse), Life Technologies/Invitrogen Alexa 488 donkey anti-mouse, Alexa 568 donkey anti-rabbit, Alexa 647 donkey anti-goat. Protease inhibitors used for cell extraction and membrane preparation, leupeptin, Pefabloc, E-64, antipain, aprotinin, bestatin, and calpain inhibitors I and II, were purchased from A.G. Scientific, Inc. Rhodamine, rhodamine green, Latrunculin B, cytochalasin D, FITC-phalloidin, TRITC-phalloidin, blebbistatin, were purchased from Sigma. MLCK inhibitor ML-7 (4310) was purchased from Tocris. Alexa 647-phalloidin and Alexa 350-phalloidin were purchased from ThermoFisher/LifeTechnologies. Stock solutions of latrunculin B (5 mM), cytochalasin D (2 mM), blebbistatin (5 mM), ML-7 (10 mM), phalloidin were prepared in dimethylsulfoxide.

### DNA constructs

Synaptopodin A was synthesized by Genscript using the coding sequence of human synaptopodin (NCBI Reference Sequence: NP_009217.3) and was subcloned by Genscript into HINDIII and Xho1 sites of the blasticidin-selectable mammalian expression vector pcDNA6 myc-HisA (Invitrogen) containing an N-terminal sstFRET (Meng and Sachs, 2011) and C-terminal myc and His tags.

sstFRET consists of a spectrin repeat flanked by cerulean and Venus fluorescent proteins (tccgtgagcaagggcgaggagctgttcaccggggtggtgcccatcctggtcgagctggacggcgacgtaaacggccacaagttcagcgtgtccggcgagggcgagggcgatgccacctacggcaagctgaccctgaagctgatctgcaccaccggcaagctgcccgtgccctggcccaccctcgtgaccaccctgggctacggcctgcagtgcttcgcccgctaccccgaccacatgaagcagcacgacttcttcaagtccgccatgcccgaaggctacgtccaggagcgcaccatcttcttcaaggacgacggcaactacaagacccgcgccgaggtgaagttcgagggcgacaccctggtgaaccgcatcgagctgaagggcatcgacttcaaggaggacggcaacatcctggggcacaagctggagtacaactacaacagccacaacgtctatatcaccgccgacaagcagaagaacggcatcaaggccaacttcaagatccgccacaacatcgaggacggcggcgtgcagctcgccgaccactaccagcagaacacccccatcggcgacggccccgtgctgctgcccgacaaccactacctgagctaccagtccaaactgagcaaagaccccaacgagaagcgcgatcacatggtcctgctggagttcgtgaccgccgccgggatcactctcggcatggacgagctgtacaagtccggactcagatctggaggcttccacagagatgctgatgaaaccaaagaatggattgaagagaagaatcaagctctaaacacagacaattatggacatgatctcgccagtgtccaggccctgcaacgcaagcatgagggcttcgagagggaccttgcggctctcggtgacaaggtaaactcccttggtgaaacagcagagcgcctgatccagtcccatcccgagtcagcagaagacctgcaggaaaagtgcacagagttaaaccaggcctggagcagcctggggaaacgtgcagatcagcgcaaggcaaagggaggcgtgaattccatggtgagcaagggcgaggagctgttcaccggggtggtgcccatcctggtcgagctggacggcgacgtaaacggccacaagttcagcgtgtccggcgagggcgagggcgatgccacctacggcaagctgaccctgaagttcatctgcaccaccggcaagctgcccgtgccctggcccaccctcgtgaccaccctgacctggggcgtgcagtgcttcgcccgctaccccgaccacatgaagcagcacgacttcttcaagtccgccatgcccgaaggctacgtccaggagcgcaccatcttcttcaaggacgacggcaactacaagacccgcgccgaggtgaagttcgagggcgacaccctggtgaaccgcatcgagctgaagggcatcgacttcaaggaggacggcaacatcctggggcacaagctggagtacaacgccatcagcgacaacgtctatatcaccgccgacaagcagaagaacggcatcaaggccaacttcaagatccgccacaacatcgaggacggcagcgtgcagctcgccgaccactaccagcagaacacccccatcggcgacggccccgtgctgctgcccgacaaccactacctgagcacccagtccaagctgagcaaagaccccaacgagaagcgcgatcacatggtcctgctggagttcgtgaccgccgccgggatcactctcggcatggacgagctgtacaag).

ShRNA for canine synaptopodin cloned into puromycin-selectable pRS mammalian expression vector (Origene) has previously been described (Kannan and Tang, 2015). Plasmids for alpha-actinin-4 FRET-based tension sensors were synthesized and sequenced by Genscript. Alpha-actinin-4 sstFRET408 was constructed by inserting the sstFRET module between 408aa and 409aa of human alpha-actinin-4 (XP_016882820.1), between the first and second spectrin repeats, and cloned into the blasticidin selectable pUNO expression vector (Invivogen). Alpha-actinin-4 sstFRET522 was constructed by inserting the sstFRET module between 408aa and 409aa of human alpha-actinin-4 (XP_016882820.1), between the second and third spectrin repeats, and cloned into pUNO. Plasmids for alpha-actinin-1 FRET-based tension sensors were kindly provided by Fanjie Meng & Frederick Sachs, Physiology and Biophysics Department, SUNY at Baffalo (Meng and Sachs, 2011). Alpha-actinin-1-M-sstFRET containing the sstFRET module inserted between 300aa and 301aa within the first spectrin repeat of human alpha-actinin-1 (P12814.2) was in the neomyocin selectable pcDNA3.1 expression vector (Invitrogen/ThermoFisher). Actinin-C-sstFRET containing the sstFRET module added to the C-terminus of human alpha-actinin-1 (P12814.2) was in pcDNA3.1. Plasmids for mEmerald-occludin (54211), mCherry-ZO-1 (55165), and mCherry-vinculin (55159) were purchased from Addgene. Plasmids for bacterial expression of recombinant alpha-actinin-4 was generated from PCR products using the coding sequence of human alpha-actinin-4 (NP_004915.2) kindly provided by Martin Pollak. Alpha-actinin-4 full length (1-911aa), actin binding domain (1-270aa), and spectrin repeats/EF hands (271-911aa) were subcloned into EcoR1 and Xho1 sites in the kanamycin-selectable bacterial expression vector pET30a+ containing an N-terminal 6His tag (EMD4Biosciences) and the ampicillin-selectable bacterial expression vector pMAL4cx with N-terminal maltose-binding protein.

### Cell culture & transfection

HUVEC (OCS-100-013), C2bbE1 (CRL-2102), and T84 (CCL-248) cells were purchased from ATTC. Madin-Darby canine kidney (MDCK) cells were originally from Kai Simons lab (EMBL, Germany) and a gift from Barry Gumbiner (University of Washington). The cells have been authenticated by staining of E-cadherin and ZO-1 using antibodies that only recognize the canine proteins, RR1 for E-cadherin and R40.76 for ZO-1. The cells are free from mycoplasma contamination as determined by the original source. MDCK cells were maintained in MEM/Earle’s balanced salt solution supplemented with 25 mM HEPES and 5% fetal bovine serum (FBS) in a 37-degree incubator in the presence of 5% CO2. Media was changed every 2-3 days. Cell stocks for parental and transfected cells are stored in freezing media (normal growth media, 20% FBS, 10% DMSO) in liquid nitrogen. MDCK cells were grown on tissue culture plastic dishes. For live-cell imaging, wound healing, and immunofluorescence, cells were plated on No.2 VistaVision cover glasses (VWR). For hydraulic chamber experiments and immunofluorescence, cells were plated on 12 mm polyester Transwell-clear with 0.4 um pores (Corning). Confluent monolayer of MDCK epithelial cells polarized on semi-permeable Transwell support for >2 weeks reaches a steady-state that generate junctions with strong cell-cell adhesion.

For transfection, cells were incubated in a 1:1 mixture of DNA/Polyjet DNA transfection reagent for 5-18 hours according to the manufacturer’s protocol (SignaGen Laboratories). Expression of plasmid DNA was selected using G418, puromycin, or blasticidin for 10-14 days. Stable expressing clonal cell lines were obtained as published (Kannan and Tang, 2015). Briefly, antibiotic resistant clonal cell lines were expanded and assessed for knockdown efficiency and protein expression by western blot and immunofluorescence. Clonal cell lines with homogeneous knockdown phenotype were used for a second round of transfection with ShRNA. Secondary clonal cell lines were expanded and assessed for knockdown efficiency by western blot and immunofluorescence. Clonal cell lines with knockdown efficiency of >80% were used for this study.

For live-cell imaging of alpha-actinin-4 sstFRET522, alpha-actinin-1-M-sstFRET and alpha-actinin-1-C-sstFRET, and synaptopodin-venus cell lines were expanded and assessed for expression using live-cell microscopy. For mixed population of cells expressing alpha-actinin-4 sstFRET408, antibiotic resistant cells were pooled and used for live-cell imaging 14-21 days post-transfection or frozen until use.

### Drug treatment and Wound closure assay

Warmed normal growth media with or without cytochalasin D (at a final concentration of 0.2 uM), latrunculin (at a final concentration of 5 uM) or blebbistatin (at a final concentration of 60 uM) were added to MDCK confluent monolayers one hour before fixation for immunofluorescence staining. For wound studies, MDCK confluent monolayers were grow on non-permeable glass coverslips for >3-4 weeks. After dome formation, monolayers were rinsed 3 times with calcium-free phosphate-buffered saline and incubated with calcium-free serum-free MEM media at 37 degrees for 10-15 min until the cells forming the domes were detached from the monolayers. Immediately after that, the calcium-free media was removed and fresh pre-warmed normal growth media with or without cytochalasin D (at a final concentration of 0.2 uM), latrunculin (at a final concentration of 5 uM) or blebbistatin (at a final concentration of 60 uM) was gently added to the dishes containing the coverslips with attached MDCK monolayers that now have holes. Cell monolayers were incubated at 37-degree incubator for wound closure to proceed. At one hour after initiation of wound healing, cell monolayers on coverslips were processed immediately for immunofluorescence.

### Staining and immunofluorescence of cell

Cells grown on Transwell-Clear (Corning) were used in localization studies. For immunofluorescence, cells were rinsed twice in 150 mM NaCl/2 mM CaCl_2_/2 mM MgCl_2_/20 mM HEPES, pH 7.8 and fixed in 1% formaldehyde/150 mM NaCl/2 mM CaCl_2_/2 mM MgCl_2_/20 mM HEPES, pH 7.8 at 4°C for 1 hour. The reaction was quenched with Tris in quenching buffer (0.05% Triton X-100/50 mM Tris/100 mM NaCl/20 mM HEPES, pH 8.0) for 3 hours. The Transwell with fixed cells were rinsed in immunofluorescence staining buffer (0.05% Triton X-100/100 mM NaCl/20 mM HEPES, pH 7.5), the cells were incubated with primary antibodies (1 ug/ml) in staining buffer overnight. After rinsing in staining buffer three times, the cells were incubated in secondary antibodies in staining buffer for 90 min. Then, the cells were rinsed three times in staining buffer and incubated with fluorescently labeled phalloidin or Hoechst 33528 for 60 min. Finally, the cells were rinsed three times in staining buffer and post-stain fixed with 1% formaldehyde in staining buffer for 3 hours. Transwell filters were excised using a razor blade and mounted with No.1 glass coverslips on glass slides with ProLong Glass antifade (Invitrogen). Stained cells with antifade were allowed to cure for 48 hours before used for imaging.

### Image acquisition of fixed cell

Images were collected in 200-nm steps using Axio Imager.Z2m microscope equipped with Apotome.2 (Carl Zeiss) and X-cite 120 LED (Lumen Dynamics). For Optical Sectioning Structured Illumination Microscopy (OS-SIM), 7 phases/images were collected per each constructed image using either alpha Plan-Apochromat 100x/NA1.46 Oil DIC M27 or Plan-Apochromat 40X/NA1.3 Oil DIC M27 objectives (Carl Zeiss) and a 4K ORCA-Flash4.0 V2 digital CMOS camera with 6.5 um x 6.5 um pixel size (Hamamatsu Photonics). Wide-field optical z images were deconvolved using the Zen2 pro deconvolution module. Low magnification wide field images were collected using Plan Apochromat 20X/0.8 objectives (Zeiss). Images for figures 1A, 1B, and 7A were collected using a 2K Optimos CMOS camera with 6.5 um x 6.5 um pixel size (Qimaging, Photometrics). Composite images were generated using ImageJ (NIH) and Zen (Carl Zeiss) softwares. For figure generation, images were cropped, contrasted, and scaled using Photoshop software (Adobe) before importing into Illustrator (Adobe).

### Image analysis

All images were corrected for chromatic shift on the X, Y, Z-axes for each fluorescence channel before were used for analysis. Quantitation of actin (phalloidin) and junction proteins (immunofluorescence intensity) was performed in ImageJ (NIH) or Zen (Carl Zeiss) using unprocessed original single optical z-slice images. A defined area was used to compare the signal intensity of actin (phalloidin) and immunofluorescence of junctional proteins. All measured intensities were subtracted from background signal (an area with no cells or within the cytoplasm) before used for statistical analyses and calculation of intensity ratios. All experiments had been repeated at least three times. At least 6 data set from each experiment were collected.

All analyses were performed using KaleidaGraph software (Synergy). Each junctional region is outlined using a rectangle box tool and each contractomere is outlined using a circle tool. The mean pixel intensity of each defined junctional region is used for comparison of junction localization of individual protein. For calculation of Pearson’s correlation coefficient R, intensities of individual pixel within the defined junctional region were used and each pixel corresponds to 45 x 45 nm of the imaged sample. All p-values were calculated using non-paired student t-tests. Linear regression was fitted using original data points. Line intensity graphs were generated in Excel (Microsoft) using pixel intensities from original images. For measurement of junctional length, distance between cell vertices, cell perimeter, wound perimeter and area, a free-drawing tool and a line tool were used to trace outlines and draw straight lines.

### Live-cell imaging setup

For live-cell imaging, glass coverslips were soaked in 100% ethanol, and sterilized under UV for 60 min. Sterilized coverslips were coated with 20 ug/ml collagen IV in phosphate buffered saline for 60 min and used immediately for plating of cells. Cells grown on collagen-coated glass coverslips were mounted up-side-down onto an in-house fabricated polycarbonate chamber with a deep well for media for long-term imaging. Imaging was performed in FluoroBrite/DMEM (Gibco) media containing 10% fetal bovine serum and 10 mM HEPES, pH 7.5. Sample temperature was maintained at 35 degrees Celsius on a heated stage and an objective heater (PeCon). Images were collected using ORCA-Flash4.0 (Hamamatsu Photonics) or Optimos (Qimaging) mounted onto Axio Imager.Z2m (zeiss) with X-cite 120 LED (Lumen Dynamics). Representative movies are presented in the paper. Additional movies will be provided upon request.

### Live-cell imaging of alpha-actinin FRET and synaptopodin

Alpha-actinin tension sensors were imaged using the Gemini dual-view system (Hamamatsu Photonics) equipped with excitation filter for Cerulean fluorescence protein (CeFP) and emission filters for Cerulean fluorescence protein (CeFP) and Venus fluorescence protein (VFP). Simultaneous acquisition of images for CeFP and VFP emission were obtained using a Plan-Apochromat 40x/NA1.3 Oil DIC M27 objective (Zeiss). Junctional intensities of CeFP and VFP were used to calculate the FRET index (EmVFP)/(EmCeFP + EmVFP), which is shown as FRET/(CeFP+FRET). Briefly, the CeFP and VFP channels were overlay on top of each other using a macro written within the Zen2 software (Carl Zeiss). Each junctional region is outlined manually with a rectangle drawing tool in the Zen2 imaging tool (Zeiss). Contractomeres were circled with 9-pixel diameter and 160 nm per pixel resolution. The intensities of the junctional signal were measured and subtracted from background signal (an area with no cells or within the cytoplasm) before used for the calculation of the FRET index.

For movie generation, individual images of cropped cells were imported into Image J (NIH) or QuickTime (Apple) to generate avi or mpeg files. Composite images were generated using ImageJ (NIH) or Photoshop (Adobe). Individual cell diameter and junction for each image were measured using image J (NIH) or Zen (Carl Zeiss). For drug treatments, cells were treated for one hour before imaging in pre-warmed normal growth media with or without cytochalasin D (at a final concentration of 0.2 uM), ML-7 (at a final concentration of 10 uM) or Latrunculin B (at a final concentration of 5 uM).

### Immunoprecipitation

Stable clonal cell line expressing alpha-actinin tension sensors were plated at confluent density and allowed to grow for 2 days. Cell were extracted with 1% TX-100 in 150 mM NaCl, 20 mM HEPES, pH 7.5 in the presence of protease inhibitors (10 ug/ml Leupeptin, 1mg/ml Pefabloc, 10 ug/ml E-64, 2 ug/ml antipain, 2 ug/ml aprotinin,50 ug/ml bestatin, 20 ug/ml calpain inhibitors I & 10 ug/ml calpain inhibitor II). Insoluble materials were pelleted by spinning total cell extract for 10 min at 1000xg. Supernatants were incubated overnight with anti-Venus antibodies (MABE1906, EMD Millipore) or anti-alpha-actinin-4 antibodies (sc-393495, Santa Cruz Biotechnology). The antibodies-cell lysates were then incubated with Protein G Dynabeads (10003D, ThermoFisher Scientific) for 2 hours. The beads were washed 4 times in extraction buffer and twice in 5 mM HEPES, pH7.5. The bound fractions were solubilized in SDS-PAGE sample buffer as described above.

### Western blots

For comparison of young and mature monolayers, confluent monolayers of cells were trypsinized and replated at high confluent density (10^7^ cells per 10 cm). Cells were allowed to form cell-cell interactions for 2 days (young) or 7 days (mature). Total cell lysates were obtained by solubilizing the cells in SDS-PAGE sample buffer containing 25 mM dithiothreitol, 2% SDS, 50 mM Tris-Cl, 5% glycerol, pH 8.8 and protease inhibitors (10 ug/ml Leupeptin, 1mg/ml Pefabloc, 10 ug/ml E-64, 2 ug/ml antipain, 2 ug/ml aprotinin,50 ug/ml bestatin, 20 ug/ml calpain inhibitors I & 10 ug/ml calpain inhibitor II). For western blot of phosphorylated myosin light chain, cell extraction was carried out as above with the addition of a phosphatase inhibitor cocktail (Simple Stop 1 #GB-450) purchased from Gold Biotechnology, USA. Biorad DC detergent compatible protein assay was used to determine total protein concentration in cell lysates. Equal protein amounts of cell lysates were used for comparison of junctional protein. SDS-PAGE was performed using 8-16% gradient gel and transferred to nitrocellulose paper using the Transblot (BioRad). Western blots were carried out using iBind with 1 ug/ml primary antibodies and 1 ug/ml secondary antibodies following iBind protocol (Invitrogen/ThermoFisher).

### Active Rho Detection

Rho activity was assayed using the Active Rho Detection Kit (Cell Signaling Technology #8820). Briefly, confluent monolayers of MDCK and synaptopodin knockdown cells were lyzed and the supernatants were used for immunoprecipitation using GST-Rhotekin-Rho-binding domain. The immunoprecipitated protein complexes were eluted in SDS sample buffer and ran on an SDS-PAGE gel. Rho levels in Rhotekin immunoprecipitations were detected by performing a western blot using anti-Rho antibodies.

### Recombinant protein expression, purification, and fluorophore labeling

For expression of recombinant proteins, cDNA plasmids were transformed into Rosetta DE3 E. Coli cells containing tRNA’s for “universal” translation” (Novagen) maintained under chloramphenicol. For recombinant 6-His tagged *α*-actinin-4, actin binding domain, and spectrin repeats/EF hands, protein expressions were induced with 500 uM Isopropyl β-D-1-thiogalactopyranoside for 8 hours at 25^0^ C in LB containing chloramphenicol. Cells were centrifuged at 6000xg for 15 min and resuspended in 20 mM NaCl, 20 mM HEPES, pH7.8 in the presence of 5 mg/ml lysozyme. After one freeze-thaw cycle, lysed cells were centrifuged at 100,000xg for 30 min. Supernatant was loaded onto a nickel column (Qiagen). The column was washed with 20 bed volumes of 500 mM NaCl, 25 mM Imidazole, and 20 mM HEPES, pH 7.8. The recombinant proteins were eluted with 10 bed volumes of 500 mM NaCl, 500 mM Immidazole, 20 mM HEPES, pH7.8. Eluted proteins were concentrated using Centricon filters (Millipore, Inc) and purified by gel filtration using Superdex 200 in 150 NaCl, 20 mM HEPES, 10 mM *β*-mercaptoethanol. Proteins were either frozen or used immediately for labeling. Recombinant alpha-actinin-4 proteins were labeled on cysteine using maleimide-activated Oregon Green at a ratio of 5 fluorophores for every *α*-actinin at room temperature for one hour. Labelled proteins were separated from free dyes by gel filtration using Superdex 200 in 150 NaCl, 20 mM HEPES, 10 mM *β*-mercaptoethanol.

For expression of heterodimeric alpha-actinin-4 consists of one monomer of full-length and one monomer of spectrin-repeats/EF hands, plasmid pet30a+ with full-length alpha-actinin-4 and plasmid pMAL-4cx with spectrin repeats/EF hands were transformed together into Rosetta DE3 cells. Cells expressing both constructs were selected with ampicillin and kanamycin and maintained in LB containing 10 mM glucose to suppress the expression of MBP-fusion protein. To induce protein expression, 200 uM Isopropyl β-D-1-thiogalactopyranoside was added to the LB growth media for 8 hours at 25 degrees. Cells were centrifuged at 6000xg for 15 min and resuspended in 20 mM NaCl, 20 mM HEPES, pH7.8 in the presence of 5 mg/ml lysozyme. After one freeze-thaw cycle, lysed cells were centrifuged at 100,000xg for 30 min. Supernatant was loaded onto an amylose column 3 times (NED), washed with 20 bed volumes of 100 mM NaCl, 20 mM HEPES, pH7.8, and eluted with 10 mM maltose in 100 mM NaCl, 20 mM HEPES, pH7.8. The eluted fraction was loaded onto a nickel column (Qiagen). The column was washed with 20 bed volumes of 500 mM NaCl, 25 mM Imidazole, and 20 mM HEPES, pH 7.8. The recombinant proteins were eluted with 10 bed volumes of 500 mM NaCl, 500 mM Immidazole, 20 mM HEPES, pH7.8. Eluted proteins were concentrated using Centricon filters (Millipore, Inc) and purified by gel filtration using Superdex 200 in 150 NaCl, 20 mM HEPES, 10 mM *β*-mercaptoethanol.

Actin was purified from rabbit skeletal muscle as described (Brieher et al., 2004). Actin was labelled on lysine residues using two NHS activated rhodamine for every actin molecule for one hour at room temperature as described (Brieher et al., 2006). After ’labeling, actin filaments were pelleted at 100,000 X g for 30 minutes, resuspended in G buffer and dialyzed exhaustively against G buffer (0.2 mM ATP, 0.2 mM CaCl_2_, 5 mM Tris-HCl, pH8, 1 mM *β*-mercaptoethanol). This procedure typically labeled actin to 50-80%. Aliquots of proteins were snap-freeze in liquid nitrogen and stored at -80 degrees until use. Protein concentrations were determined by Bradford assay (BioRad).

### Purification of junction-enriched membrane

Junction-enriched membranes were prepared as described (Tang and Brieher, 2012). Briefly, frozen rat livers (Pelfrez) were thawed in 2 volumes of 10 mM HEPES, pH8.5/10 mM DTT. Protease inhibitors (see above) were added to the thawed livers and the livers were blended in a Waring blender (5 x 15 sec). The liver slush was filtered through 4 layers of cheesecloth to obtain the total liver homogenate. Total liver homogenate was centrifuged at 1000xg for 30 min. The pellet was homogenized in 10 mM HEPES, pH8.5/10 mM DTT in a Dounce homogenizer and centrifuged at 100xg for 30 min. The supernatant was collected and centrifuged at 1000xg for 30 min. The membrane pellet contains the majority of actin assembly activity and is frozen at -80 degrees before further purification immediately before use. The day of the experiment, membranes were thawed on ice, diluted 1:1 with 10 mM HEPES, pH 8.5 supplemented with 10 mM DTT and homogenized through a 25G needle. The homogenates were spun through a 20% sucrose pad for 10 minutes at 16,000xg. The supernatant was discarded, and the pellet were resuspended with 10 mM HEPES, pH 8.5 supplemented with 10 mM DTT. The homogenate was spun through a 20% sucrose pad for 15 min at 1000xg. The pellet was discarded, and the supernatant was spun through a 20% sucrose pad for 15 min at 16,000xg. The membrane pellet contains junction-enriched plasma membrane fragments as determined by western blots and immunostaining (Tang and Brieher, 2012).

### Membrane actin assembly and immunofluorescence

Actin assembly reactions were performed in actin polymerization buffer (50 mM KCl, 2 mM EGTA, 2 mM MgCl_2_, and 100 mM HEPES, pH 7.8) supplemented with 2 mM buffered ATP, pH 8. A standard 20-µL reaction consists of ∼15 µg of total proteins from the purified junctional membrane fraction and 1 µM rhodamine or Oregon green-labeled monomeric actin. Actin polymerization was allowed to carry out at room temperature for 30-60 min.

For reconstitution assays, purified membranes were stripped with high salt (500 mM NaCl, 2 mM MgCl_2_, 2 mM EGTA, 20 mM HEPES, pH 7.8, and 10 mM DTT), TX-100 or CHAPS (in 50 mM NaCl, 2 mM MgCl_2_, 2 mM EGTA, 20 mM HEPES, pH 7.8, and 10 mM DTT) on ice for 1 hour. Stripped membranes were collected by centrifugation through a 20% sucrose cushion at 10,000 *g* for 10 min. Purified proteins (labelled and unlabeled α-actinin-4) were allowed to bind to stripped membranes for 1 hour at room temperature. Unbound proteins were removed by spinning membranes through a 20%w/w sucrose cushion at 10,000 *g* for 10 min. The final reconstitution reaction consists of ∼8 µg of total protein from stripped membranes, 0.5 µM fluorescently labeled monomeric actin, and purified proteins (α-actinin-4) and was carried out at room temperature for 30 min.

For immunofluorescence of membranes, actin assembly assay was performed, and membranes were collected by centrifugation through a 20% sucrose cushion at 10,000 *g* for 10 min. The purified membranes with incorporated actin were incubation with primary antibodies in the presence of 0.1 % TX-100 in actin assay buffer for 2 hours. The membranes were spun through a 20% sucrose cushion and resuspended in 0.1% TX-100 in assay buffer. The membranes were incubated with secondary antibodies for 2 hours, spun through a 20% sucrose cushion, and resuspended in 0.1% TX-100 in assay buffer. The membranes were mounted on a glass slide and covered with a No.1.5 glass coverslip before imaging. Images were obtained using a Plan-Apochromat 63x/NA1.4 Oil DIC M27 objective (Carl Zeiss) attached to an Axio Imager (Carl Zeiss) equipped with ORCA-ER CCD camera with 6.45 um X 6.45 um pixel size (Hamamatsu Photonics) and the Colibri Illumination System (Carl Zeiss).

### Electron Microscopy

For visualization of actin with the junctional complex on membrane sheet, actin assembly reaction was performed using rhodamine-labeled monomeric actin at a final concentration of 1 µM. Actin polymerization was allowed to carry out at room temperature for 60 min. The reaction was purified by spinning through a 20% sucrose cushion. The membrane with incorporated actin was resuspended in 10 uL of actin polymerization buffer. 4 uL of the resuspended membranes was used for wide-field light microscopy as described above. 5 uL of the resuspended membranes was processed for electron microscopy analysis. The resuspended membranes were allowed to attach onto glow-discharged carbon-coated grids for 10 min. Unbound membranes were removed by washing the EM grids three times with assembly buffer. The membranes were negatively stained with 2% uranyl acetate and excess stain was remove immediately. The grids were allowed to air dry for 10 min and stored until image under an electron microscope. Images were collected with a JEOL 2100EX at 120 kV using a 2K × 2K CCD camera (UltraScan; Gatan, Inc.). For figure generation, images were cropped, contrasted, and scaled using Photoshop software (Adobe) before importing into Illustrator (Adobe).

## Data Availability

All data are available from the corresponding author.

## Acknowledgements

We thank Nivetha Kannan, Cameron Shahnazi, Kevin Huang, Kyle Sherman, Bobby Knier, Rafael Anorga for help with protein gels, molecular biology, generation and maintenance of cell lines, and molecular biology. Electron microscopy was carried out in part in the Frederick Seitz Materials Research Laboratory Central Facilities, University of Illinois. The pressure chamber was built by the mechanical engineering machine shop at the University of Illinois, Urbana-Champaign. This work is funded by the National Institute of Health (R01-DK098398 to Vivian W Tang and R01-GM106106 to William M Brieher). The authors declare no competing financial interests.

## Author Contribution

Timothy Morris generated synaptopodin reagents for cell experiments. Eva Sue generated alpha-actinin-4 cell lines for FRET experiments. Caleb Geniesse purified junctional membranes and generated alpha-actinin-4 proteins for in vitro experiments. William M Brieher provided actin reagents and funding. Vivian M Tang performed experiments, analyzed data, wrote manuscript and provided funding.

---

**Figure S1.**
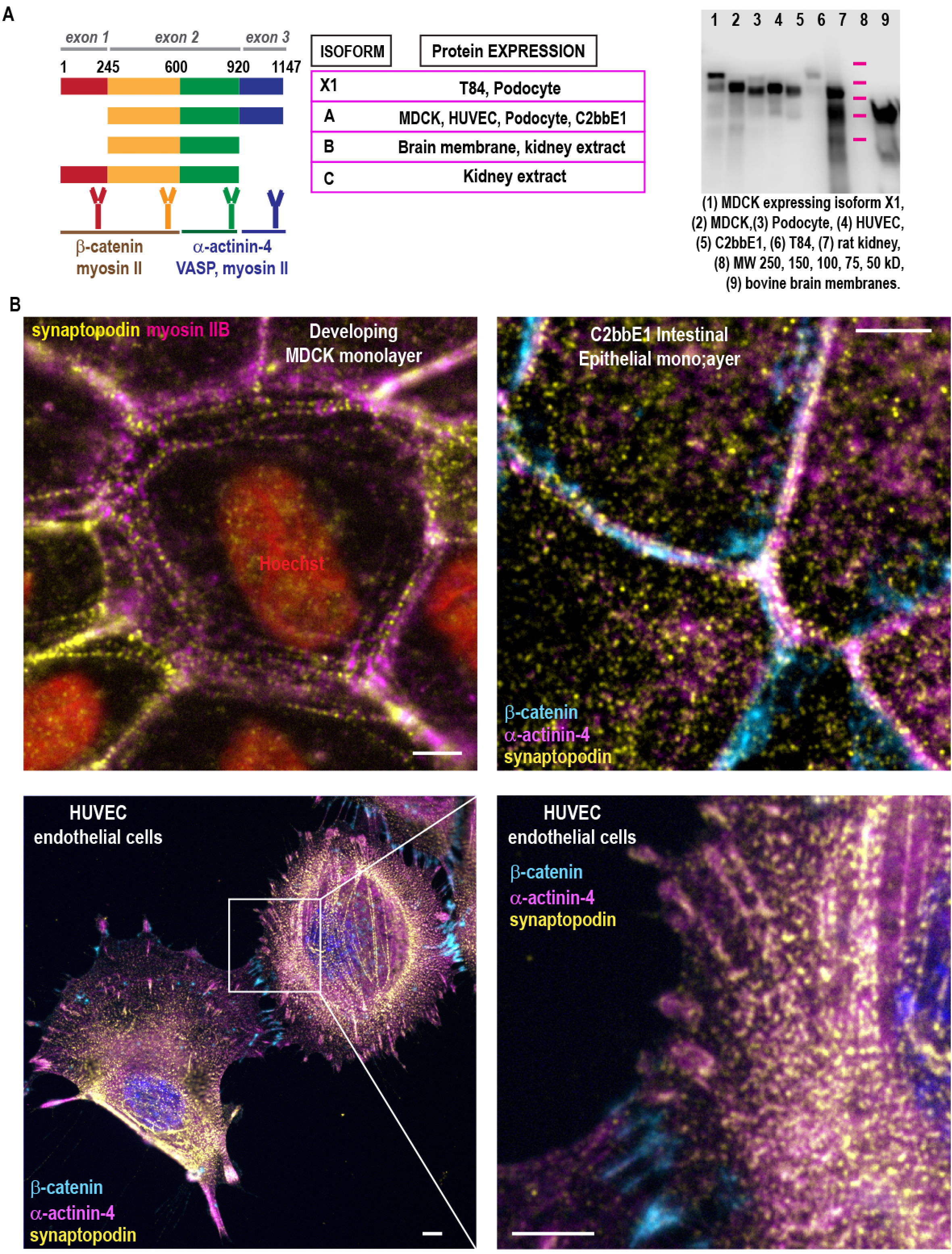
Synaptopodin is localized to junctions and stress fibers of epithelial and endothelial cells. (A) Synaptopodin is encoded by 3 exons, resulting in 4 splice variants. We raised antibodies to each spliced region to characterize the expression of synaptopodin isoforms in mammalian tissues and cells. Western blot using antibodies against region encoded by exon 2, thus recognizing all synaptopodin isoforms. Table summarized results from western blot studies using our newly generated antibodies (see methods section). (B) Immunofluorescence of MDCK kidney tubule epithelial cells, C2bbE2 intestinal epithelial cells, HUVEC endothelial cells. Scale bars are 2 microns.

**Figure S2.**
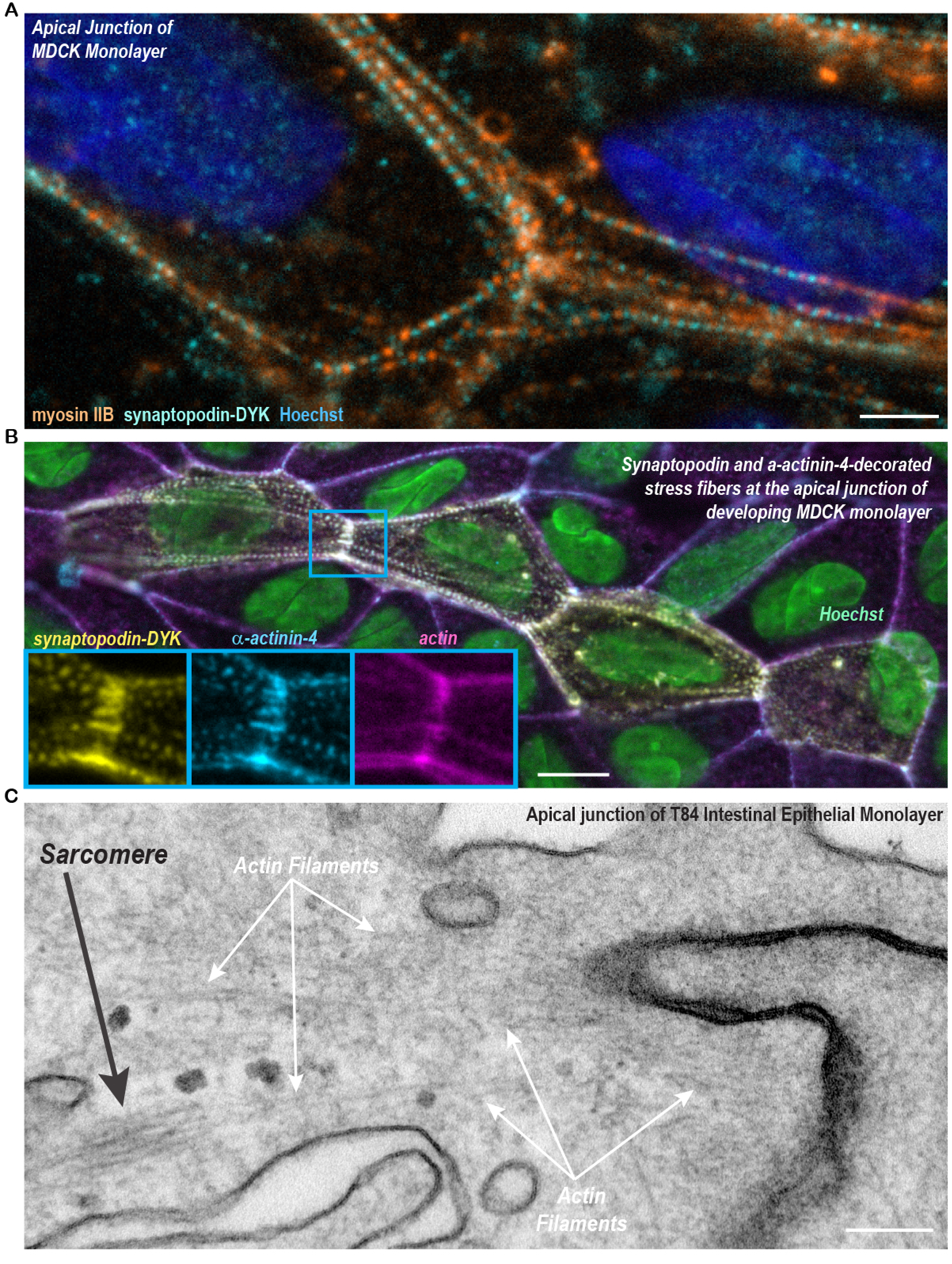
Apical stress fibers insert into the apical junctions of epithelial cells. (A) Immunofluorescence showing alternating arrangement of synaptopodin and myosin IIB forming sarcomeric-like repeats. Scale bar is 2 microns. (B) Immunofluorescence showing apical stress fibers inserted into cell-cell adhesions to connect multiple cells. Insets show synaptopodin and alpha-actinin-4 overlap on apical stress fibers. Scale bar is 5 microns. (C) Thin-section transmission electron microscopy showing actin bundle connected to the apical junction on one end and a sarcomeric structure on the opposite end. Scale bar is 200 nm.

**Figure S3.**
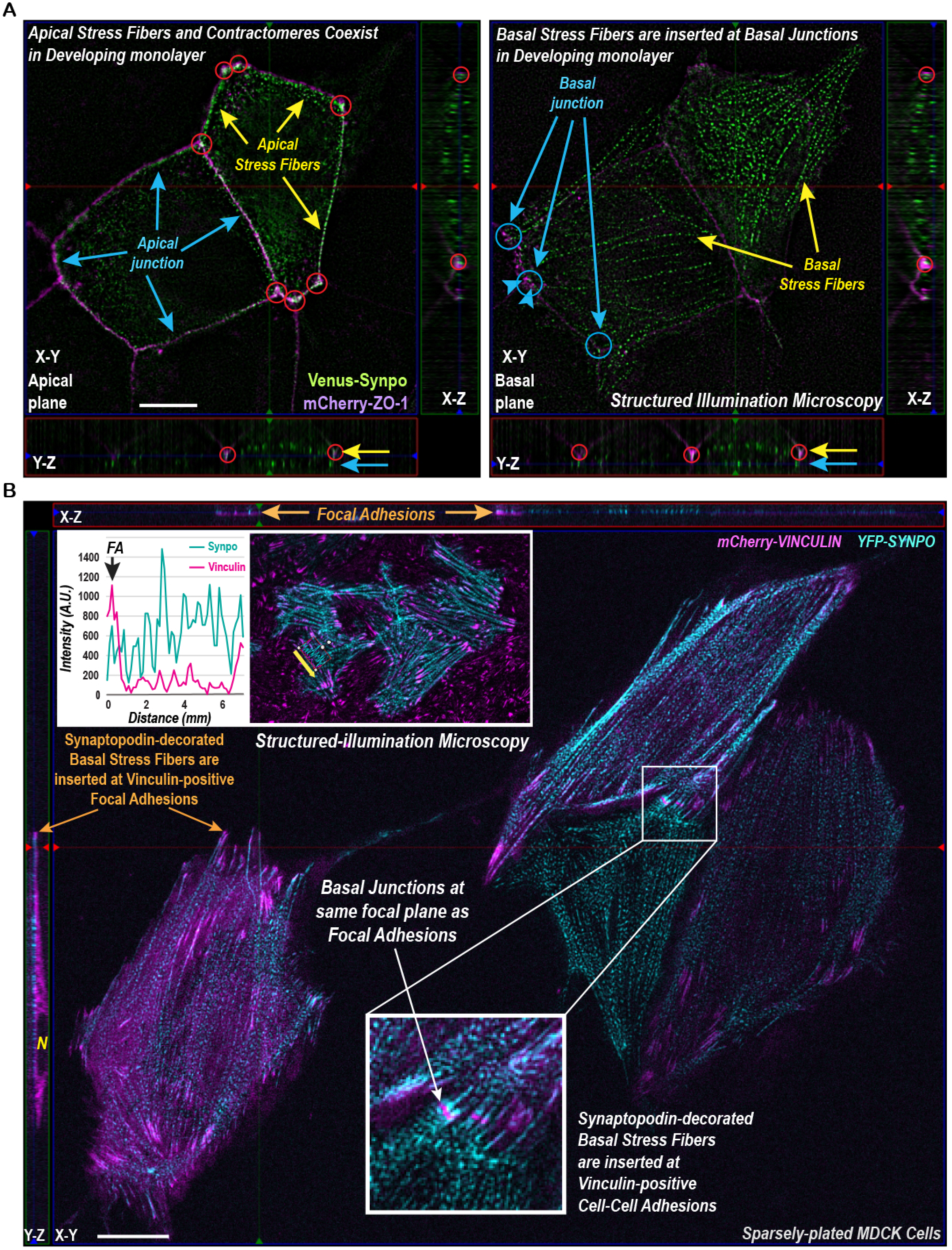
Vinculin marks the basal junction whereas ZO-1 marks both apical and basal junctions. (A) Structured-illumination microscopy of synaptopodin and ZO-1. A z-stack is shown in X-Y, Y-Z, and X-Y axes. Left panel shows apical stress fibers and contractomeres on the apical plane. Right panel shows basal stress fibers inserted at basal junctions. Attachment sites for synaptopodin apical stress fibers are circled. Arrows on Y-Z axis point to apical and basal planes. Scale bar is 5 microns. (B) Live-cell structured-illumination microscopy of synaptopodin and vinculin. Vinculin marks the basal junction where basal stress fibers are inserted. Inset shows synaptopodin stress fibers inserted at vinculin-decorated focal adhesions. Graph shows a line scan of the arrow along a basal stress fiber in the inset. Synaptopodin has periodic organization at the stress fiber. Scale bar is 5 microns.

**Figure S4.**
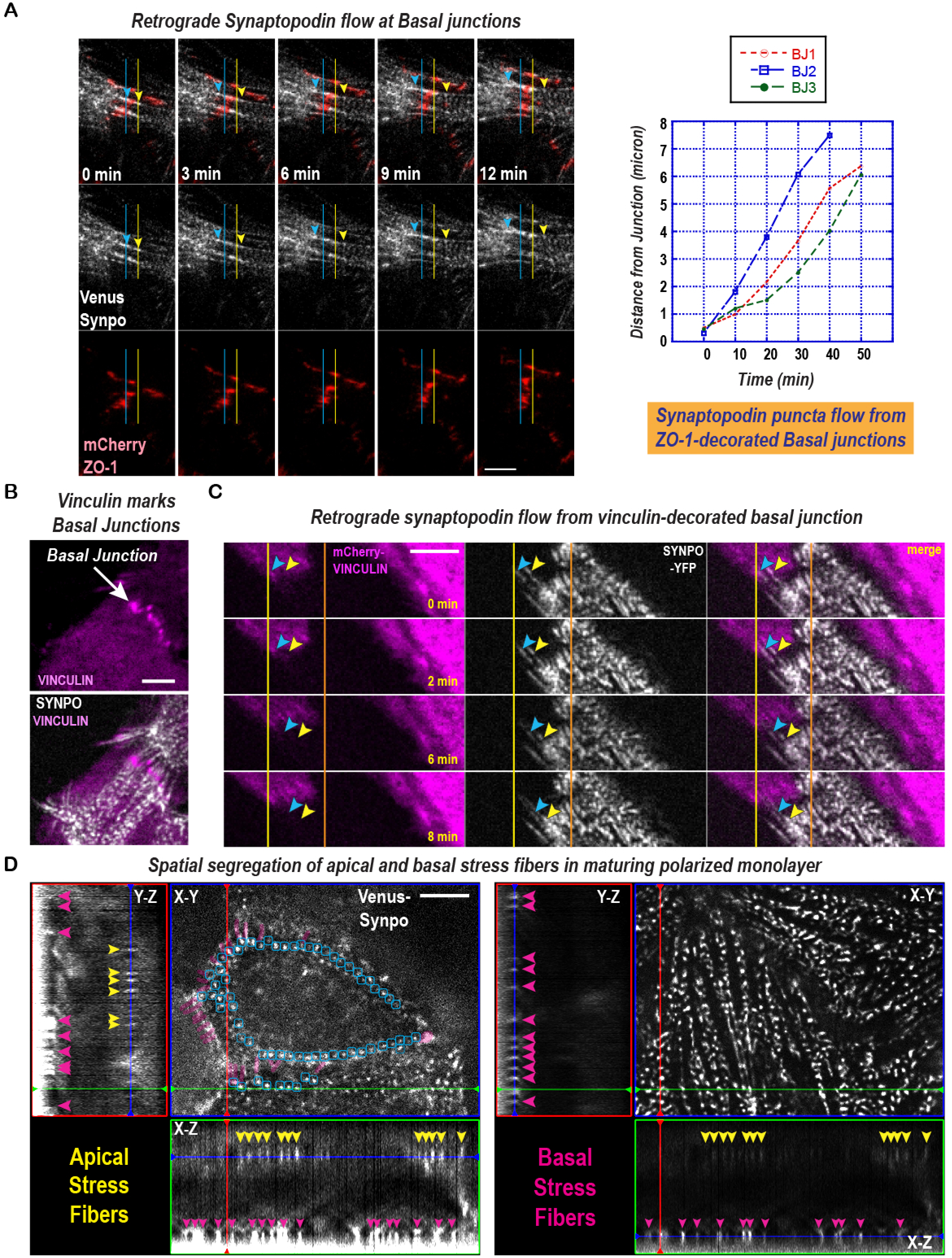
Retrograde synaptopodin flow from the basal junction. (A) Frames taken from a time-lapse of synaptopodin and ZO-1. Synaptopodin retrograde flow on opposite sides of a basal junction. Arrowheads track the flow of synaptopodin puncta in 2 cells. Graph shows rate of retrograde flow by tracking synaptopodin puncta. Scale bar is 1 micron. (B) Structured-illumination live-microscopy of synaptopodin and vinculin showing insertion of stress fibers from 2 cells at basal junctions. Scale bar is 2 microns. (C) Frames taken from a time-lapse movie of synaptopodin and vinculin. One cell is expressing synaptopodin to show retrograde synaptopodin flow from basal junctions marked by the neighboring cell. Arrowheads track the flow of synaptopodin puncta. Scale bar is 1 micron. (D) Live imaging of synaptopodin. Left panel shows the apical plane and right panel shows the basal plane of the cell; X-Y, Y-Z, and X-Z views are shown. Synaptopodin linkers at the apical junction is highlighted in purple. The repeated and aligned synaptopodin densities are circled in blue. Yellow arrowheads mark synaptopodin at apical stress fibers in X-Z and Y-Z views. Pink arrowheads mark synaptopodin at basal stress fibers in Y-Z view. The periodic spacing of synaptopodin densities are seen in both apical and basal stress fibers. Scale bar is 2 microns.

**Figure S5.**
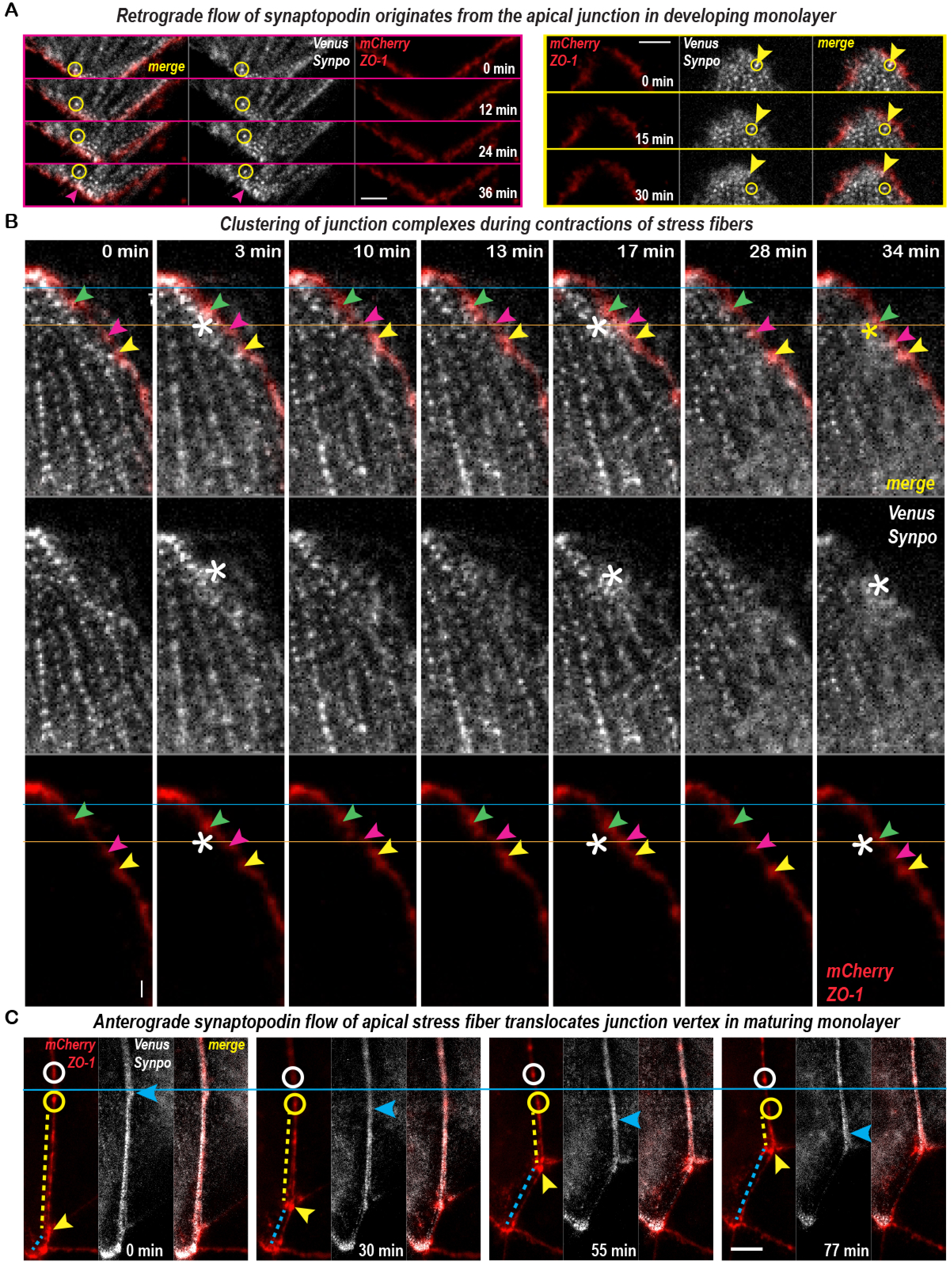
Live imaging of synaptopodin showing retrograde flow during early development, contraction during maturation, and anterograde flow towards junctional vertex during late stage of maturation. (A) Frames from time-lapse of synaptopodin and ZO-1 in cells with developing junctions. Retrograde synaptopodin flow originating from the apical junction. Circles track single synaptopodin densities as they flow inward from the apical junction into the medial-apical region. Scale bars are 1 micron. (B) Frames from time-lapse of synaptopodin and ZO-1 in cells with maturing junctions. ZO-1 densities, marked by arrowheads, are temporarily clustered when synaptopodin stress fiber contract (asterisk). (C) Frames from time-lapse of synaptopodin and ZO-1. Anterograde synaptopodin flow towards the junction vertex is associated with the movement of junctional vertex into the synaptopodin flow. Circles track synaptopodin densities flowing into the junction vertex. Arrowhead track the movement of junction vertex. Scale bar is 500 nm.

**Figure S6.**
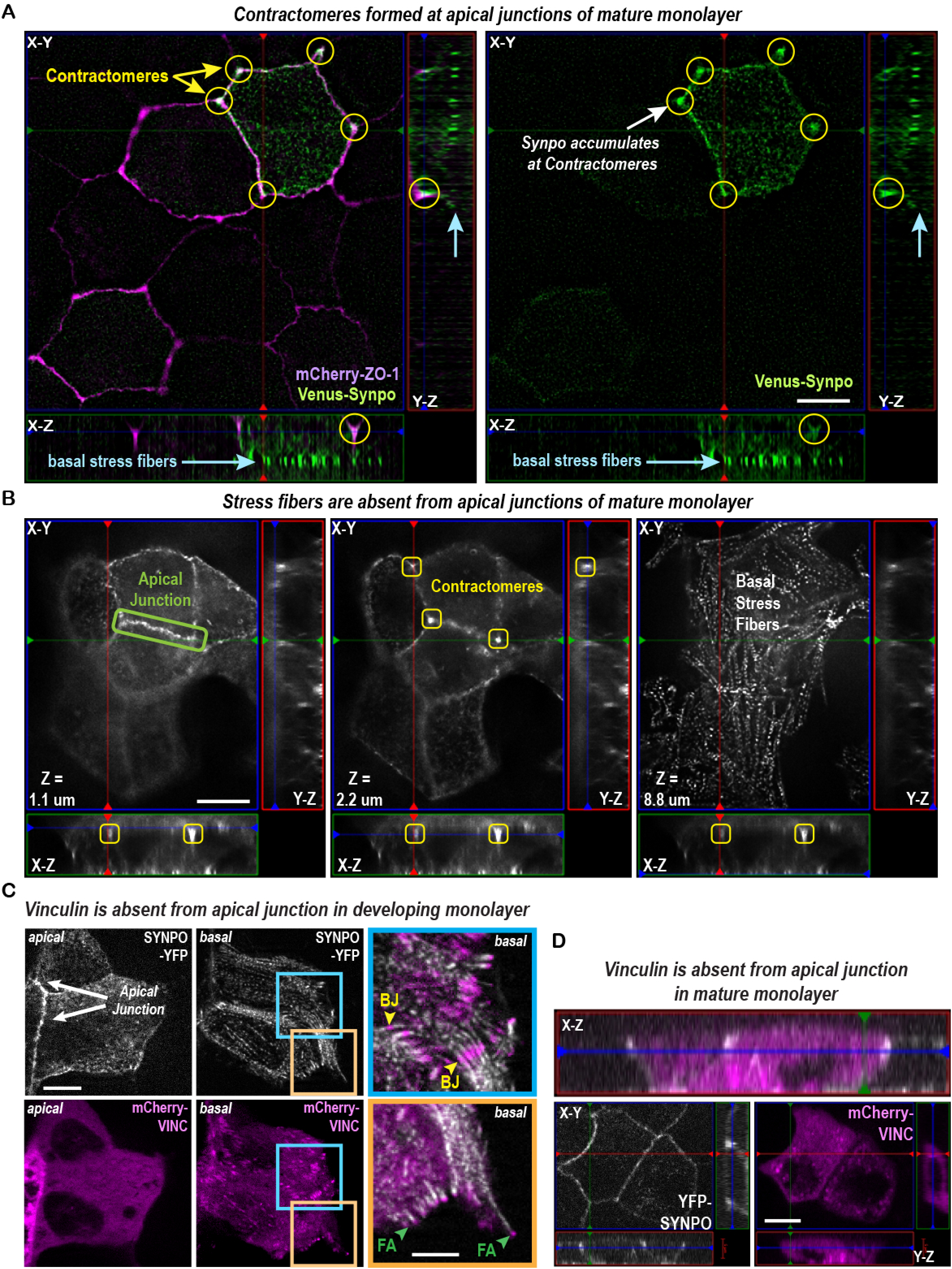
Contractomeres formation at late stage of junction maturation. (A) Structured-illumination microscopy of synaptopodin and ZO-1. A z-stack is shown in X-Y, Y-Z, and X-Y axes. Left panel shows contractomeres on the apical plane. Right panel shows basal stress fibers. Arrows on Y-Z axis point to the basal plane. Scale bar is 5 microns. (B) Disintegration of periodic synaptopodin organization and formation of contractomeres at the apical junction in mature monolayer. Left and middle panels show the top and apical junction , and the right panel shows the basal palne; X-Y, Y-Z, and X-Z views are shown. Basal stress fibers remain intact despite disassembly of apical stress fibers. Contractomeres are squared. Scale bar is 5 microns. (C) Live-cell Structured-illuminated microscopy of synaptopodin and vinculin. Vinculin is absent from the apical junction in developing monolayer. Blue and orange boxes are higher magnification showing basal junctions and focal adhesions on the same basal focal plane. Scale bars are 2 microns. (D) Vinculin is absent from the apical junction in mature monolayer. A z-stack is shown in X-Y, X-Z, and Y-Z views. Scale bar is 5 microns.

**Figure S7.**
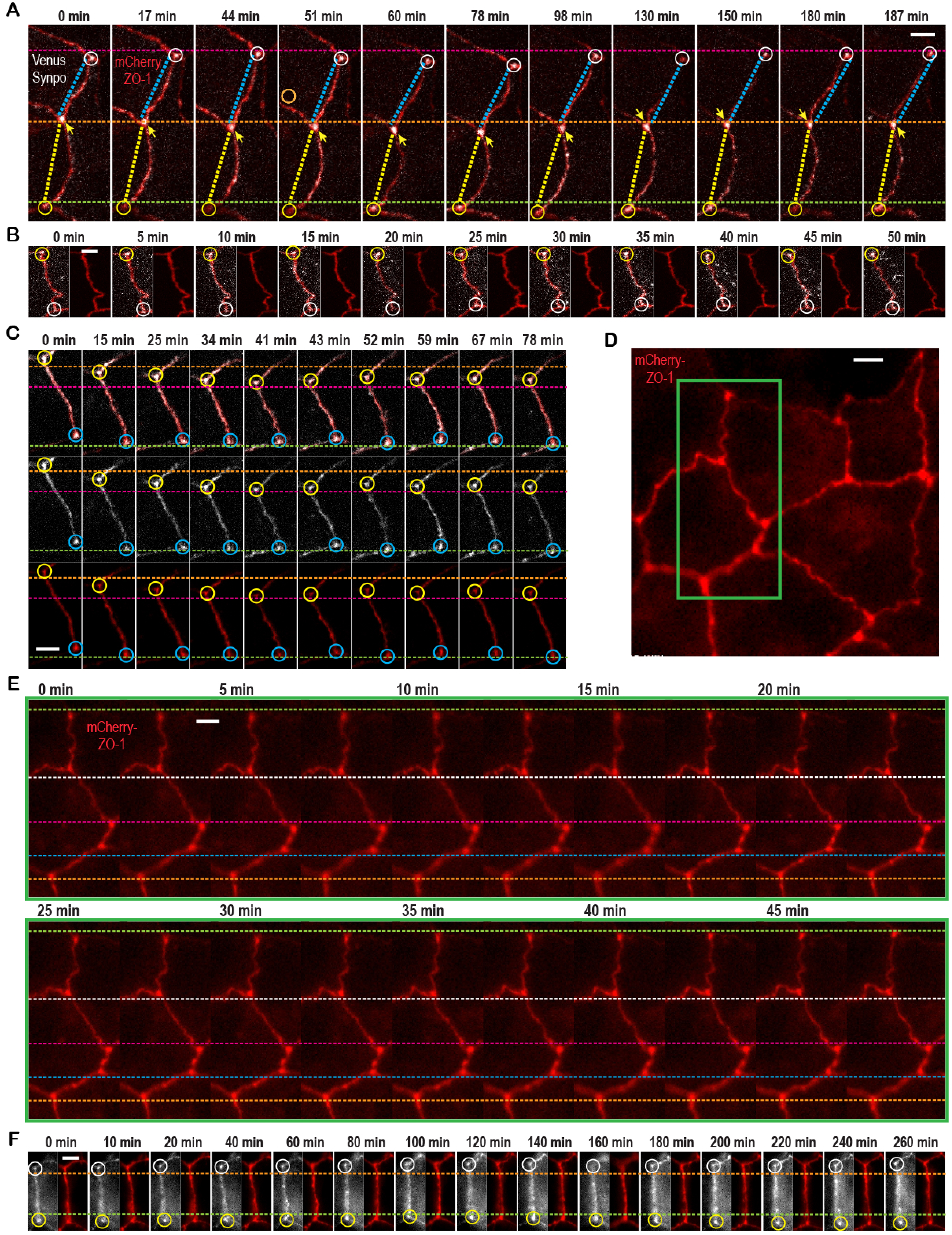
Contractomeres are stationary in mature monolayer. (A-F) Frames from time-lapse movies showing oscillation of contractomeres with no net motility or change in junctional lengths. Scale bars are 1 micron.

**Figure S8.**
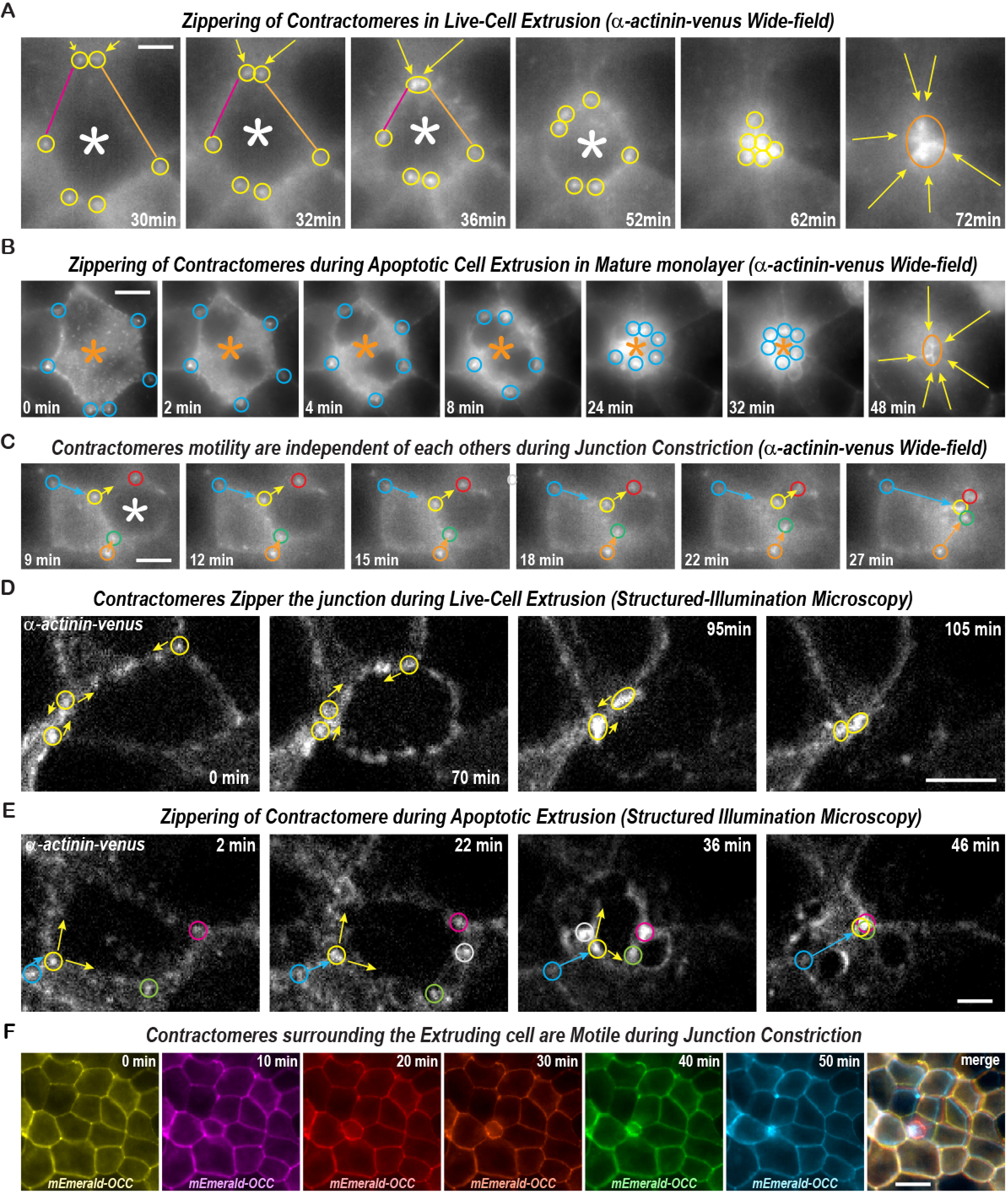
Contractomere motility during cell extrusion in mature monolayer. (A) Frames from time-lapse movie showing gliding of contractomere (yellow circles) glides toward each other to constrict the apical junction during live-cell extrusion (white asterisk). Yellow arrows mark the paths of contractomere movement. Orange circle shows the location of contractomeres after constriction is completed. Scale bar is 2 microns. (B) Frames from time-lapse movie showing gliding of contractomere (blue circles) towards each other to constrict the apical junction during apoptotic cell extrusion (orange asterisk). Yellow arrows mark the paths of contractomere movement. Orange circle shows the location of contractomeres after constrictions is completed. Scale bar is 10 microns. (C) Frames from time-lapse movie showing gliding of contractomeres surrounding the extruding cell (white asterisk). Contractomeres circled in blue and orange are immobile while contractomeres next to the extruding cell (circled in yellow, green, and red) are mobile. Movements of contractomeres next to the extruding cell (yellow, red, green circled) shorten the apical junction surrounding the extruding cell (white asterisk) and lengthen the apical junction in the neighboring cells (blue and orange arrows). Scale bar is 5 microns. (D) Frames from time-lapse structured-illumination microscopy showing gliding of contractomeres (circles) to constrict the apical junction (white asterisk). Scale bar is 5 microns. (E) Frames from Structured-illumination microscopy showing gliding of contractomeres (circles) to shorten the junction surrounding the extruding cell (yellow arrows) while lengthen the junction in the neighboring cells (blue arrow). Scale bar is 2 microns. (F) Frames from time-lapse movie showing one cell extrusion events. Merge image shows overlapping junctions except the ones immediately surrounding the extruding cells. Scale bar is 10 microns.

**Figure S9.**
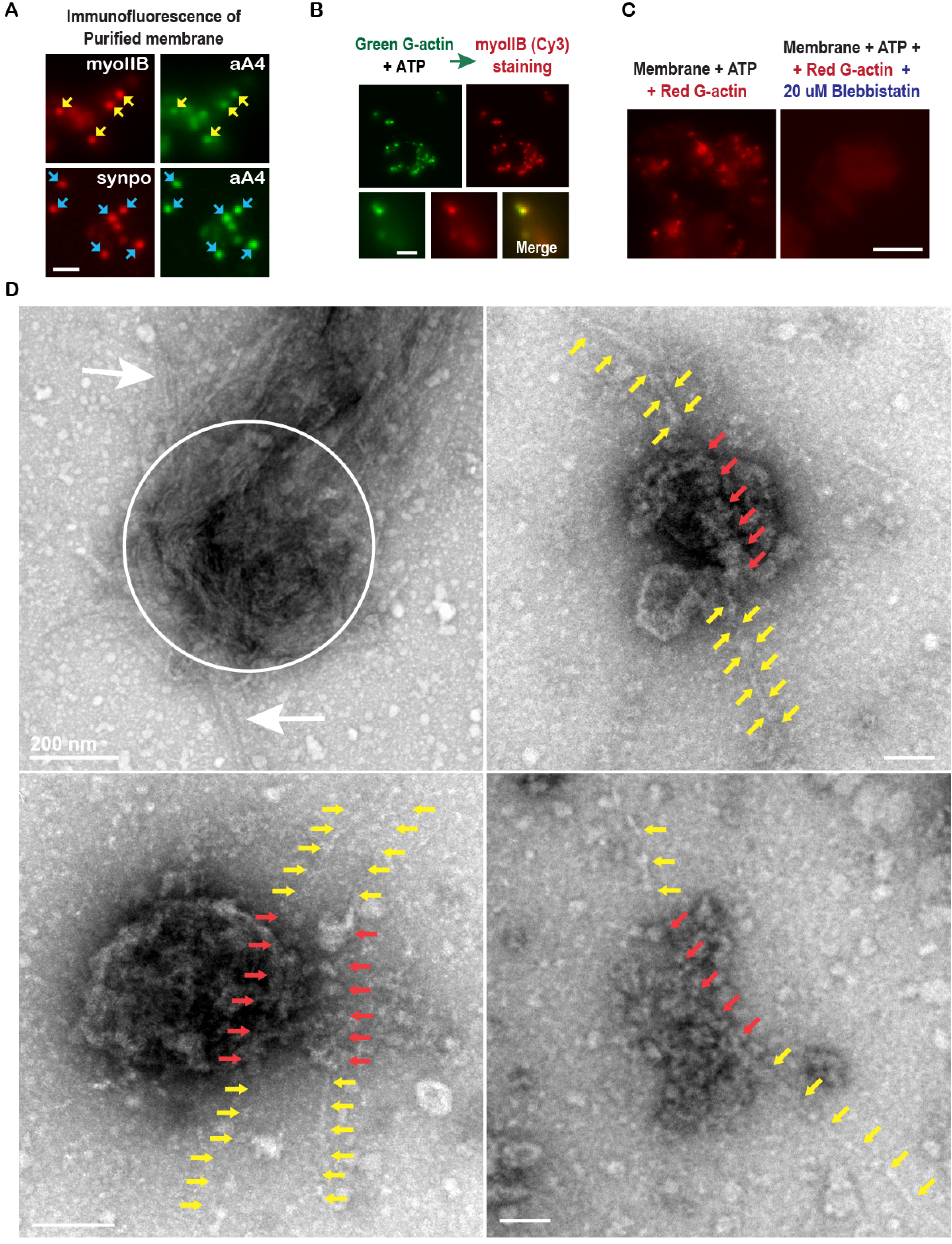
Unconventional biochemistry for actin assembly at the contractomere. (A) Immunofluorescence of junctional membranes showing myosin IIB and synaptopodin colocalize with alpha-actinin-4 on contractomeres. Scale bar is 1 micron. (B) Myosin IIB is localized to sites of actin assembly on junction membranes. Actin is initiated by adding G-actin to purified membranes in the presence of ATP (see Methods). Immunofluorescence staining for myosin IIB was performed on membranes after an actin assembly assay. Scale bar is 500 nm. (C) Actin assembly on purified membrane is blocked by the addition of blebbistatin to the actin assembly mixture (see Methods). Scale bar is 5 microns. (D) Negative-stain electron microscopy of contractomeres after actin assembly reaction (see Methods). Upper left panels show actin bundles wrap around to form a ball (circled) after an actin assembly reaction using 2 micromolar of G-actin. Single actin filaments (arrows) can be seen in associated with the actin ball. Scale bar is 200 nm. Upper right and lower left panels showing actin filaments (yellow arrows) interact with multiple globular densities (red arrows) on the contractomeres after an actin assembly reaction using 500 nM of G-actin. Scale bars are 100 nm. Lower right panel shows a contractomeric complex extracted with CHAPS detergent to remove lipids components. Multiple densities (red arrows) are interacting with an actin filament (yellow arrows). Scale bar is 50 nm.

**Figure S10.**
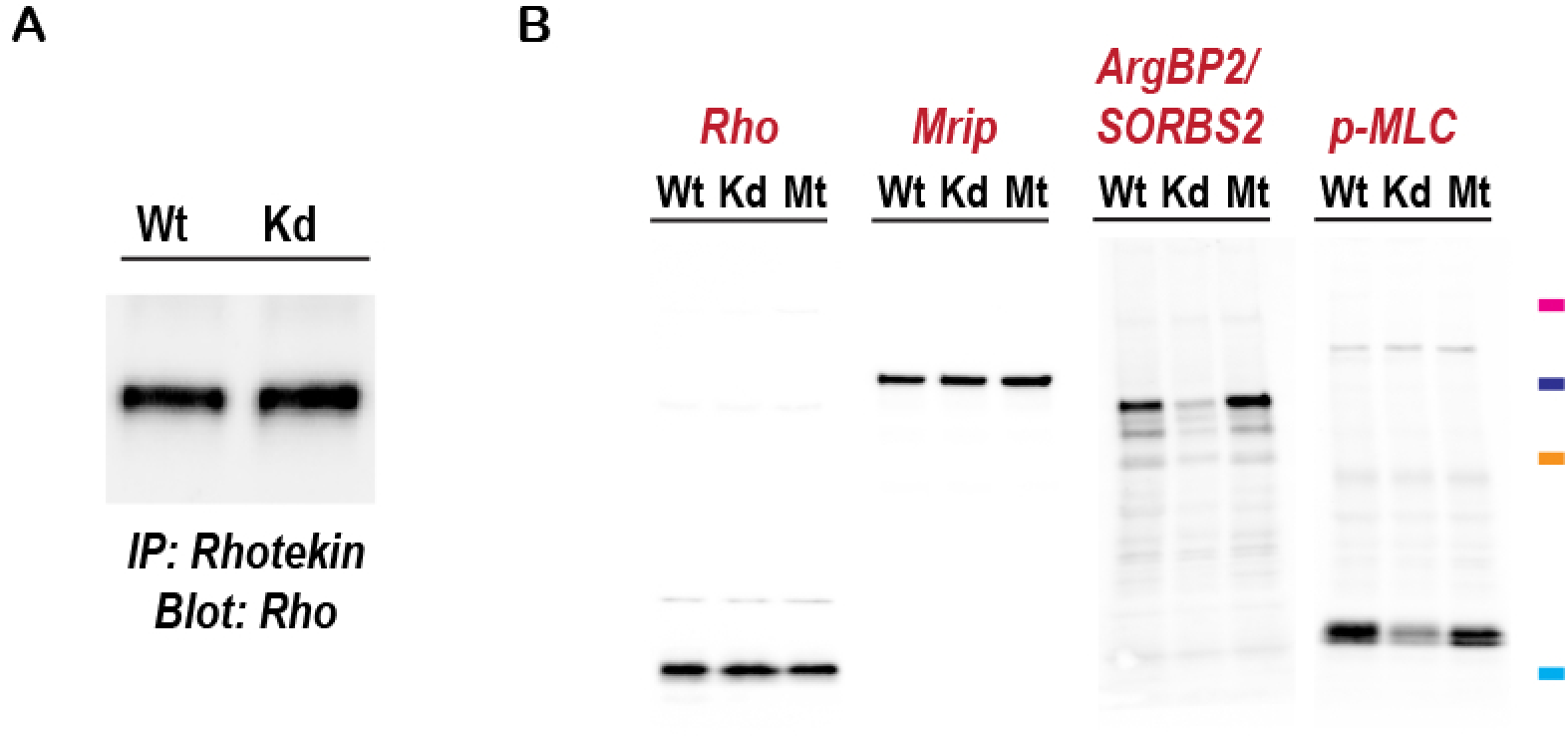
Synaptopodin knockdown reduces ArgBP2/SORBS2 levels without affecting RhoA. (A) Synaptopodin knockdown does not affect RhoA activity in MDCK cells as assessed by Rhotekin pull-down and RhoA western blot (see Methods). (B) Western blots showing decrease in ArgBP1/SORBS2 and phospho-myosin light chain levels in synaptopodin knockdown cells whereas RhoA and MRIP levels were unaffected. Markers are 100 75, 50, and 25 kD.

## Movie Legend

Movie 1. Live-cell time-lapse of Venus-synaptopodin and mCherry-ZO-1 showing contractions of apical stress fibers concomitant with junction remodeling.

Movie 2. Live-cell time-lapse of Venus-synaptopodin and mCherry-ZO-1 showing contractions of apical stress fibers concomitant with reshaping of cell boundary and junction remodeling.

Movie 3. Live-cell time-lapse of Venus-synaptopodin and mCherry-ZO-1 showing retrograde flow of synaptopodin from the basal cell junctions.

Movie 4. Live-cell time-lapse of Venus-synaptopodin and mCherry-ZO-1 showing contractions of basal stress fibers and retrograde flow of synaptopodin from the basal cell junctions.

Movie 5. Live-cell time-lapse of Venus-synaptopodin and mCherry-vinculin showing retrograde flow of synaptopodin from the basal cell junction. Venus-synaptopodin and mCherry-vinculin are expressed in 2 neighboring cells showing synaptopodin stress fibers inserted at sites of vinculin enrichment in the neighboring cell.

Movie 6. Live-cell time-lapse of Venus-synaptopodin and mCherry-ZO-1 showing retrograde flow of synaptopodin from the apical cell junction. Synaptopodin network originated from the apical junction contracts to remodel the apical junction and the apical domain.

Movie 7. Live-cell time-lapse of Venus-synaptopodin and mCherry-ZO-1 showing retrograde flow of synaptopodin from the apical cell junction. Synaptopodin network originated from the apical junction has oscillating contractile behaviors and contracts to remodel the apical junction.

Movie 8. Live-cell time-lapse of Venus-synaptopodin and mCherry-ZO-1 showing retrograde flow of synaptopodin from the apical cell junction. Synaptopodin network originated from the apical junction has oscillating contractile behaviors and contracts to remodel the apical junction.

Movie 9. Live-cell time-lapse of Venus-synaptopodin and mCherry-ZO-1 showing retrograde flow of synaptopodin from the apical cell junction. Rows of synaptopodin originated from the apical junction has oscillating contractile behaviors.

Movie 10. Live-cell time-lapse of Venus-synaptopodin and mCherry-ZO-1 showing clustering of ZO-1 complexes during contractions of synaptopodin stress fibers parallel to the apical cell junction.

Movie 11. Live-cell time-lapse of Venus-synaptopodin and mCherry-ZO-1 showing clustering of ZO-1 complexes during contractions of synaptopodin stress fibers parallel to the apical cell junction.

Movie 12. Live-cell time-lapse of Venus-synaptopodin and mCherry-ZO-1 showing clustering of ZO-1 complexes during contractions of synaptopodin stress fibers parallel to the apical cell junction.

Movie 13. Live-cell time-lapse of Venus-synaptopodin and mCherry-ZO-1 showing clustering of ZO-1 complexes during contractions of synaptopodin stress fibers parallel to the apical cell junction

Movie 14. Live-cell time-lapse of Venus-synaptopodin and mCherry-ZO-1 showing anterograde movement of synaptopodin towards the junction vertex with concomitant movement of ZO-1 complexes towards the vertex.

Movie 15. Live-cell time-lapse of Venus-synaptopodin and mCherry-ZO-1 showing anterograde movement of synaptopodin towards the junction vertex with concomitant movement of ZO-1 complexes towards the vertex.

Movie 16. Live-cell time-lapse of Venus-synaptopodin and mCherry-ZO-1 showing motility of 2 vertices towards each other.

Movie 17. Live-cell time-lapse of Venus-synaptopodin and mCherry-ZO-1 showing anterograde movement of synaptopodin concomitant with gliding of a contractomere along the junction.

Movie 18. Live-cell time-lapse of Venus-synaptopodin showing oscillating behavior of contractomeres along the junctions in maturing monolayer.

Movie 19. Live-cell time-lapse of Venus-synaptopodin showing oscillating behavior of contractomeres in mature monolayer.

Movie 20. Live-cell time-lapse of Venus-synaptopodin showing oscillating behavior of contractomeres in mature monolayer.

Movie 21. Structured-illumination live-cell time-lapse of Venus-synaptopodin and mCherry-ZO-1 showing wiggling of apical junction and oscillation of contractomeres in mature monolayer.

Movie 22. Structured-illumination live-cell time-lapse of Venus-synaptopodin and mCherry-ZO-1 showing wiggling of apical junction and oscillation of contractomeres in mature monolayer.

Movie 23. Structured-illumination live-cell time-lapse of Venus-synaptopodin and mCherry-ZO-1 showing wiggling of apical junction, oscillation of contractomeres in mature monolayer, and removal of synaptopodin along the junction during maturation.

Movie 24. Live-cell time-lapse of mCherry-ZO-1 showing wiggling of apical junction and oscillation of contractomeres in mature monolayer.

Movie 25. Live-cell time-lapse of venus-alpha-actinin-1 showing contractomere movement during apoptotic cell extrusion. Blebbing of apoptotic cell is concomitant with cell extrusion.

Movie 26. Live-cell time-lapse of venus-alpha-actinin-1 in 4 focal planes showing contractomere movement during cell extrusion.

Movie 27. Live-cell time-lapse of venus-alpha-actinin-1 showing contractomere movement during cell extrusion.

Movie 28. Structured-illumination live-cell time-lapse of Venus-alpha-actinin-1 showing contractomere movement during live-cell extrusion.

Movie 29. Structured-illumination live-cell time-lapse of Venus-alpha-actinin-1 showing contractomere movement during apoptotic cell extrusion. Blebbing of apoptotic cell is concomitant with cell extrusion.

Movie 30. Live-cell time-lapse of mEmerald-occludin showing 3 cell extrusion events in a mature monolayer.

Movie 31. Live-cell time-lapse of mEmerald-occludin showing 1 cell extrusion event in a mature monolayer.

Movie 32. Live-cell time-lapse of Venus-synaptopodin and mCherry-ZO-1 in maturing monolayer showing wiggling of apical stress fibers in one cell and contractomere motility in an extruding neighboring cell.

Movie 33. Live-cell time-lapse of mEmerald-occludin showing 3 cell extrusion events in a mature monolayer.

Movie 34. Live-cell time-lapse of mEmerald-occludin showing 1 cell extrusion event in a mature monolayer.

Movie 35. Live-cell time-lapse of mEmerald-occludin showing 1 cell extrusion event in a mature monolayer.

## Notes

### Competing Interest Statement

The authors have declared no competing interest.

